# Changes in the neuropeptide complement correlate with nervous system architectures in xenacoelomorphs

**DOI:** 10.1101/265579

**Authors:** Daniel Thiel, Mirita Franz-Wachtel, Felipe Aguilera, Andreas Hejnol

**Affiliations:** Sars International Centre for Marine Molecular Biology, University of Bergen, Thormøhlensgate 55, 5006 Bergen, Norway; Proteome Center Tübingen, University of Tübingen, Auf der Morgenstelle 15, 72076 Tübingen, Germany

## Abstract

Neuropeptides are essential neurosecretory signaling molecules common in protostomes and deuterostomes (together Nephrozoa). Not much, however, is known about the neuropeptide complement of the sister group Xenacoelomorpha. This group is comprised of the three clades *Xenoturbella*, Nemertodermatida, and Acoela, which differ strongly in their nervous system anatomy. In order to reconstruct the ancestral bilaterian neuropeptide complement and gain insights into neuropeptide evolution within Xenacoelomorpha, we analyzed transcriptomes of 13 acoels, nemertodermatids, and *Xenoturbella* species. Together with motif searches, similarity searches, mass spectrometry and phylogenetic analyses of neuropeptide precursors and neuropeptide receptors, we reconstruct the xenacoelomorph neuropeptide complement. Our comparison of xenacoelomorph GPCRs with cnidarian and nephrozoan neuropeptide receptors shows that the neuropeptide signaling diversified into at least 20 ancestral peptidergic systems in the lineage to Bilateria. We find that *Xenoturbella* species possess many of the ancestral bilaterian peptidergic systems and only a few clade-specific neuropeptides. Nemertodermatids seem to have nearly the complete complement of ancestral bilaterian systems and several novel neuropeptides. Acoels show an extensive loss of conserved bilaterian systems, but gained the highest number of novel and group-specific neuropeptides. While it is difficult to correlate the emergence of the bilaterian neuropeptide complement with the evolution of centralized nervous systems, we find a correlation between nervous system novelties and the expansion of taxon-specific neuropeptides in Xenacoelomorpha.

## Introduction

Neuropeptides are a diverse group of small signaling molecules that play a crucial role in the function of the nervous system of most metazoan animals [1-5]. Most neuropeptides are about 3-20 amino acids long and signal via conserved GPCRs (G protein-coupled receptors) [1, 2, 6, 7]. These signaling molecules do not depend on direct synaptic transmission but often transfer signals by volume transmission and act as neuro-hormones or neuro-modulators [8-14]. Studying the distribution and expression of related neuropeptides in different animal lineages is important to understand the evolution of nervous systems [15-17]. Since the short peptides themselves often do not contain sufficient phylogenetic informative sites to reveal homology or exclude convergent evolution between distantly related animals, investigations on the evolution of peptidergic systems often have to include their receptors [1, 2, 18-22]. The comparison of neuropeptide GPCRs has shown that deuterostomes and protostomes (together Nephrozoa) share at least 30 orthologous peptidergic systems, even though many of the short peptides diverged in different lineages [1, 2, 19, 22-24]. While the complement of the protostome and deuterostome peptidergic systems is very similar, the similarities to the cnidarian systems are less obvious [2, 24]. Cnidarians generally possess many neuropeptides that are encoded in multiple para-copies on their precursors (i.e., MCPs for multi-copy peptides), but knowledge about the corresponding receptors is lacking [2, 3, 5, 25, 26]. The only clear cnidarian orthologs with nephrozoan neuropeptide GPCRs seem to be the leucine-rich repeat-containing GPCRs of glycoprotein hormone related peptides and relaxin-like peptides, both of which are evolutionarily older and also exist in placozoans [2, 5]. There are a few more putative cnidarian orthologs with nephrozoan signaling systems, but their actual phylogenetic relationship is inconclusive [2, 25, 27-29]. The available data suggests that the major radiation of the various nephrozoan peptidergic systems has occured after the cnidarian-bilaterian split, at some point in the ancestral bilaterian lineage.

Although other positions have been previously proposed [30], more thorough molecular phylogenetic analyses support the Nephrozoa and Xenacoelomorpha node as the most ancestral split within Bilateria [31-33], thus confirming earlier results [34, 35]. Investigations on the peptidergic systems in Xenacoelomorpha are therefore informative for the deeper understanding of the evolution of bilaterian nervous systems. Furthermore, xenacoelomorphs display diverse neuroanatomies with clade specific differences [36-41] to which the neuropeptide complements can be related. Xenacoelomorpha comprises three major clades, *Xenoturbella*, Nemertodermatida and Acoela, with the latter two forming the clade Acoelomorpha. The nervous system of *Xenoturbella* species is often considered to reflect a more ancestral nervous system of xenacoelomorphs as it only consists of a basiepidermal, somewhat cnidarian-like nerve net without considerable condensations [38, 40, 42-45]. Nemertodermatids possess the nerve net and additional condensed basiepidermal nerve cords that can be at different locations along the dorsal-ventral axis, as well as basiepidermal anterior condensations in many species [36, 46, 47]. The acoel nervous system is considered to have several novelties such as internalized anterior brain-like structures and often multiple subepidermal pairs of longitudinal nerve cords [36-39, 42]. Some immunohistochemical studies have demonstrated reactivity of antibodies that were raised against neuropeptides of other animals, like different RFamides or SALMFamides [39, 43, 46-52]. However, not much is known about the xenacoelomorph neuropeptide repertoire except for the presence of GPCRs that are related to FMRFamide, luqin, tachykinin and neuropeptide F receptors [53]. To better understand the origin of the bilaterian neuropeptide system complement and to test for a relationship between peptidergic systems and nervous system architectures in xenacoelomorphs, we conducted a detailed bioinformatic survey for neuropeptides and neuropeptide GPCRs in transcriptomes of 13 xenacoelomorph species. Our approach included a survey for neuropeptide precursors using sequence similarity and sequence motif searches that were complemented by a mass spectrometric analysis of peptide extracts from three acoel species. In addition, we performed a survey for neuropeptide GPCRs in Xenacoelomorpha (and Ambulacraria). This nested survey, which allows comparisons between different xenacoelomorphs, but also comparisons with other bilaterian and cnidarian neuropeptide GPCRs, provides insights into the ancestral xenacoelomorph peptidergic systems, their group specific divergences and the early diversification of today’s bilaterian neuropeptide signaling systems.

## Material and Methods

### Sequence data

We investigated assembled transcriptomes of 13 Xenacoelomorpha species from publicly available sequencing data: *Childia submaculatum* (Acoela), *Convolutriloba macropyga* (Acoela), *Diopisthoporus gymnopharyngeus* (Acoela), *Diopisthoporus longitubus* (Acoela), *Eumecynostomum macrobursalium* (Acoela), *Hofstenia miamia* (Acoela), *Isodiametra pulchra* (Acoela), *Ascoparia* sp. (Nemertodermatida), *Meara stichopi* (Nemertodermatida), *Nemertoderma westbladi* (Nemertodermatida), *Sterreria* sp. (Nemertodermatida), *Xenoturbella bocki* (Xenoturbella), *Xenoturbella profunda* (Xenoturbella). Most transcriptomes were made from several whole adults and should thus include neuronal expression (see Supplementary Table 1 for details). To obtain reference and query sequences, we collected neuropeptide-precursor and neuropeptide receptor sequences from previous publications [1, 2, 54-60] and public databases (i.e., NCBI and UniProt). We aimed for a broad sampling across Bilateria and covered different clades of chordates (i.e., Craniata, Cephalochordata, and Urochordata), ambulacrarians (i.e, Echinodermata and Hemichordata) ecdysozoans (i.e., Nematoda and Arthropoda) and spiralians (i.e., Mollusca and Annelida). To increase the amount of ambulacrarian sequences in our phylogenetic analyses, we included additional echinoderm and hemichordate transcriptomes and genome-derived proteomes in our GPCR survey: *Astrotoma agssizii* (Accession: SRR1695485), *Labidiaster annulatus* (Accession: SRR1695480, SRR1695481), *Leptosynapta clarki* (Accession: SRR1695478), *Saccoglossus mereschkowskii* (Accession: SRR1695461) *Ptychodera flava* (OIST Molecular Genetics Unit; https://groups.oist.jp/molgenu/hemichordate-genomes), *Acanthaster planci* (OIST Marine Genomics Unit, Great Barrier Reef COTS Assembly; http://marinegenomics.oist.jp/cots/viewer/download?project_id=46).

To compare the bilaterian neuropeptide GPCRs with cnidarian neuropeptide GPCRs, we used cnidarian sequences of *Nematostella vectensis* that were previously predicted and published [25] and a diverse set of receptors that were predicted by the automated NCBI annotation pipeline from the genomes of the anthozoans *Exaiptasia pallida, Acropora digitifera* and *Orbicella faveolata*.

### Identification and analysis of neuropeptide GPCRs

GPCR sequences were clustered using CLANS2 [61] and a subset of diverse sequences from each group was used as query sequences in tBLASTn searching with an e-value cutoff of 1e-30. All resulting sequences from xenacoelomorphs and ambulacrarians were used as new query sequences for an additional search to find potential hidden orthologs [62]. We analysed all retrieved sequences in a cluster analysis using CLANS2 [61] with the standard BLASTp BLOSUM 62 scoring matrix. Sequences for phylogenetic trees were aligned with ClustalX v2.1 [63], non-conserved regions were automatically removed with TrimAl [64], and phylogenetic trees were generated with RAxML v8.2.9 and FastTree v2.1 [65] using the LG amino acid substitution model. Phylogenetic trees were visualized with FigTree v1.4.3 (http://tree.bio.ed.ac.uk/software/figtree).

### Identification of neuropeptide precursors

Preproneuropeptide sequences of related neuropeptides were compared using CLANS2 and a set of diverse sequences was used as query sequences. The reference sequences were used in tBLASTn searching using BLOSUM62 and BLOSUM45 substitution matrices with an e-value cutoff of 1. In addition to sequence similarity search, we employed a custom perl script (available upon request) that uses PROSITE pattern syntax and flat files to detect specific protein sequence motifs. We screened translated transcriptomes for the presence of recurrent cleavage and amidation sites that are commonly found in multi-copy neuropeptide precursor sequences, including x(5,200)-G-[KR]-[KR]-x(2,35)-G-[KR]-[KR]-x(2,35)-G-[KR](0,1) and x(5,200)-[KR]-[KR]-x(2,35)-[KR]-[KR]-x(2,35)-[KR]-[KR]-x(2,35)-[KR]-[KR]. For possible RFamides with alternative monobasic cleavage sites, we searched for the following protein sequence motif: x(5,200)-R-F-G-[KR]-x(2,35)-R-F-G-[KR]-x(2,35)-R-F-G-[KR](0,1). This approach gave us candidate neuropeptides that were further analysed manually for recurrent peptide sequences between the cleavage sites. All sequences from the motif and similarity search with complete 5’ ends were tested for the presence of a signal peptide with Signal-3L 2.0 [66] as well as with SignalP 4.1 [67] using a D-cutoff value of 0.34. Cleavage sites were generally predicted at dibasic sites [R/K]-[R/K] and alternative monobasic cleavage sites were predicted with the online application of NeuroPred (http://stagbeetle.animal.uiuc.edu) [68]. A C-terminal glycine residue in the predicted peptides was used as an indication for C-terminal amidation of the prior amino acid residue. Results were manually examined and the positive hits were used for a reciprocal search in all transcriptomes to identify hidden orthologs [62]. Peptide sequence logos and precursor schemes were created with CLC Main Workbench (Qiagen Bioinformatics) with amino acid color-coding according to their biochemical properties (i.e., RasMol color coding). All figures were created using Adobe Illustrator CS6 with the phylogenetic relationships of Xenacoelomorpha based on [31] and [69].

### Peptide extraction and mass spectrometry (LC-MS/MS)

We extracted peptides from 5-10 specimens of whole animals of the acoels *C. macropyga, I. pulchra* and *H. miamia* following the protocol described by [55]. Collected specimens were rinsed with distilled water, transferred into the extraction buffer (90% Methanol, 9% acetic acid, 1% distilled water), grinded with a pestle and vortexed vigorously. The suspension was centrifuged at maximum speed for 20 minutes at 4°C. The supernatant was collected, completely evaporated in a vacuum concentrator and dissolved in 200 *μ*l ultrapure water. Neuropeptide mixtures were reduced and alkylated as described before [70] and desalted with C18 StageTips [71].

LC-MS analysis was performed on an EasyLC nano-UHPLC (Thermo Scientific) coupled with a Q Exactive HF mass spectrometer (Thermo Scientific). Separations of the peptide mixtures were done as previously described [55] with slight modifications. Peptides were eluted with a 57-min segmented gradient of 10–20-50-90% HPLC solvent B (80% acetonitrile in 0.1% formic acid). The mass spectrometer was operated in the positive ion mode. Full scan was acquired in the mass range from m/z 300 to 1650 at a resolution of 120,000 followed by HCD fragmentation of the 7 most intense precursor ions. High-resolution HCD MS/MS spectra were acquired with a resolution of 60,000. The target values for the MS scan and MS/MS fragmentation were 3x 10^6^ and 10^5^ charges with a maximum fill time of 25 ms and 110 ms, respectively. Precursor ions were excluded from sequencing for 30 s after MS/MS. MS/MS on singly charged precursor ions was enabled. The acquired MS raw files were processed using the MaxQuant software suite v.1.5.2.8 [72]. Extracted peak lists were submitted to database search using the Andromeda search engine [73] to query target-decoy databases consisting of the predicted propeptides and the predicted active neuropeptides, commonly observed contaminants (285 entries), and the reversed complements of those sequences. Cleavage specificity N- and C-terminal of arginine and lysine and no enzyme definition were defined. The minimal peptide length was set to four amino acids. The initial precursor mass tolerance was set to 4.5 ppm, for fragment ions a mass tolerance of 20 ppm was used. Carbamidomethylation of cysteines was defined as fixed modification in the database search. A set of expected variable modifications was defined in the database search: oxidation of methionine, acetylation of the peptide N-terminus, amidation of the peptide C-terminus, and sulfation of tyrosine. False discovery rates were set to 1% at peptide, modification site, and protein group level, estimated by the target/decoy approach [74]. Spectra of peptides having scores below 100 were validated manually.

### Data accession

All data are available from the corresponding authors upon request.

## Results

We identified homologs of metazoan and nephrozoan neuropeptides and neuropeptide receptors, as well as potential precursor sequences of various multi-copy peptides (MCPs) that seem to be specific to xenacoelomorphs. An overview of the neuropeptides and neuropeptide GPCRs in different xenacoelomorph lineages is shown in Figure 1.

**Figure 1:**
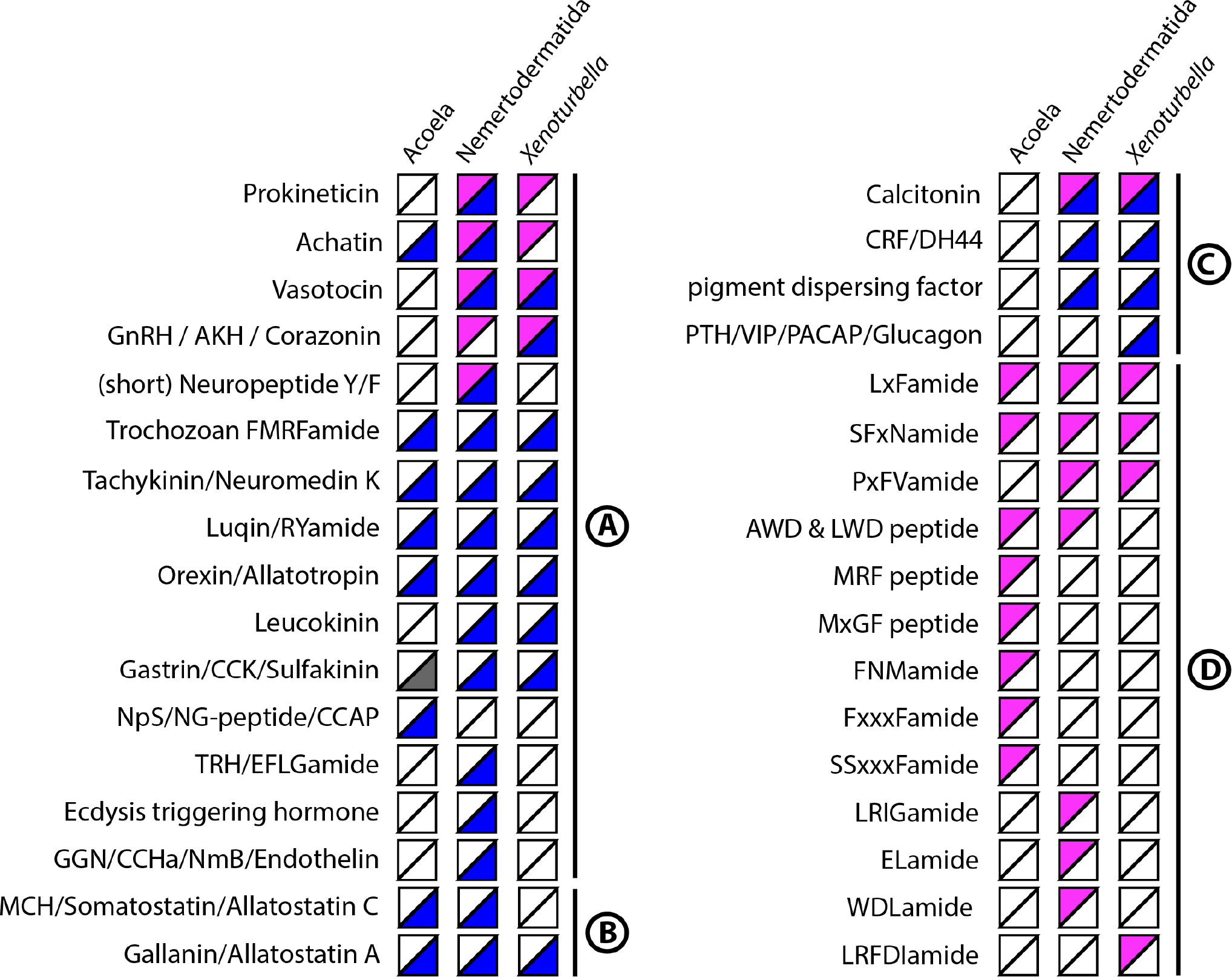
Overview of neuropeptide systems identified in Acoela, Nemertodermatida and *Xenoturbella*. Magenta upper-left half indicates at least one identified neuropeptide, blue lower-right half indicates at least one identified neuropeptide GPCR in the corresponding xenacoelomorph-clade. A rhodopsin beta-type GPCRs and potential activating ligands, B rhodopsin gamma-type GPCRs, C secretin-type GPCRs and potential activating ligands, D xenacoelomorph multi-copy peptides with orthologs in two or more species.

### Xenacoelomorph complement of plesiomorphic pre-bilaterian neuropeptides

We identified in the xenacoelomorph transcriptomes three types of neuropeptides that are present in nephrozoans as well as in non-bilaterians: glycoprotein hormone related peptides, insulin like peptides (ILPs) and prokineticin related peptides. The cnidarian-bilaterian orthology of these three peptides is supported by their highly conserved cysteine residues and partially by the presence of their receptors.

#### Glycoprotein hormone related peptides

Glycoprotein hormones or related bursicon peptides are present in all major metazoan groups [2, 5, 75]. We identified such peptides in all three major xenacoelomorph clades (Figure 2). Glycoprotein hormone alpha and beta precursors are present in both *Xenoturbella* species (i.e., *X. bocki* and *X. profunda)*. *X. bocki* contains an additional bursicon beta precursor. The nemertodermatid *N. westbladi* transcriptome contains two glycoprotein hormone alpha precursors. In acoels, we detected a homologous sequence in *I. pulchra*, which shows the cysteine arrangement of a bursicon (see [5]) but clusters in our phylogenetic analysis between glycoprotein hormone beta sequences and the bursicon alpha group, indicating a stronger sequence divergence.

**Figure 2:**
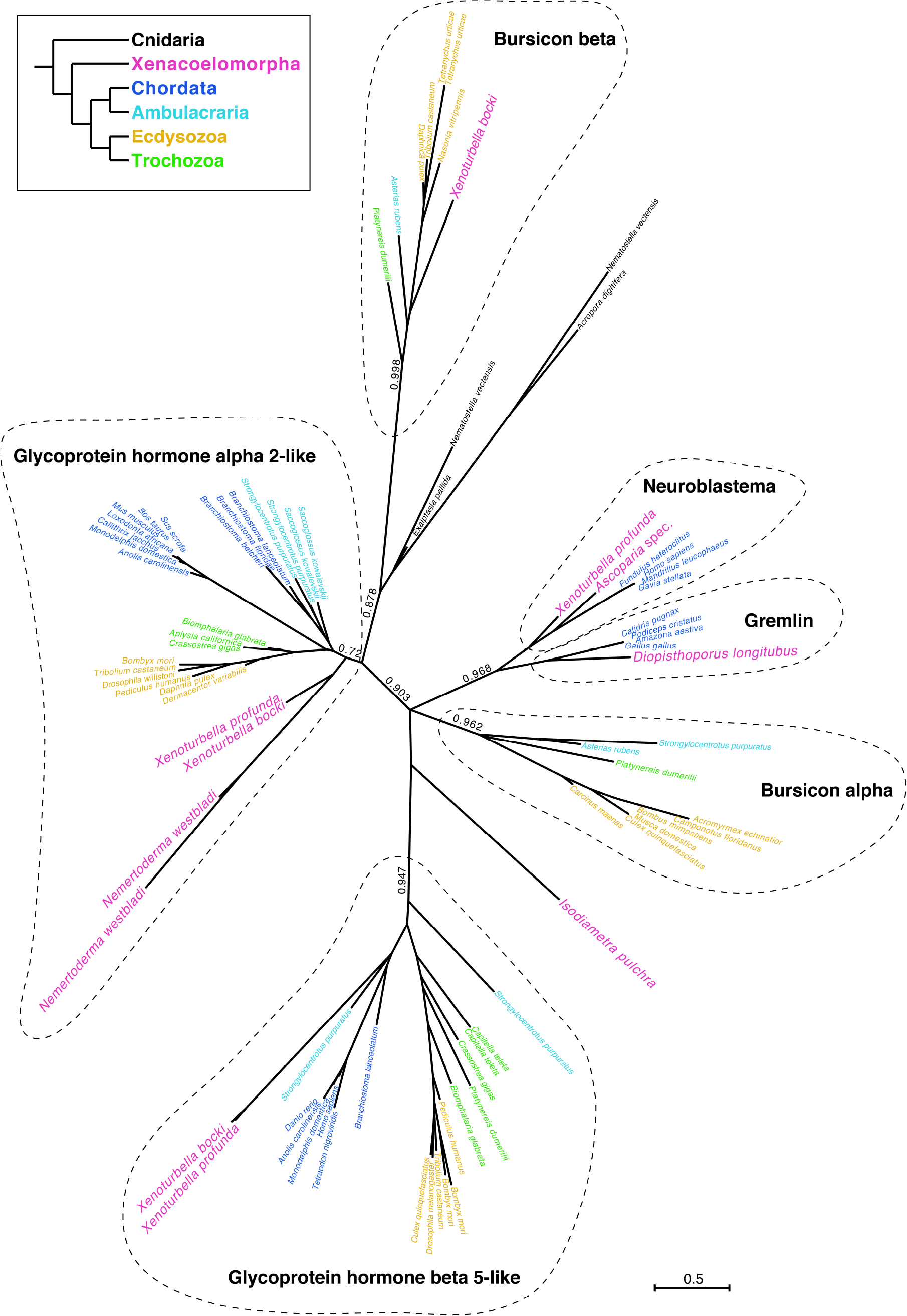
Xenacoelomorph glycoprotein hormone related peptides. Phylogenetic relationship of glycoprotein hormone and bursicon sequences. Phylogeny was calculated with FastTree v.2.1. Scale bar indicates amino acid substitution rate per site.

#### ILPs

These neuropeptides are present in all nephrozoans, cnidarians and placozoans [2, 76-79]. Many animals possess more than one ILP with an exceptional large expansion to more than 30 ILPs in the silk worm *Bombyx mori* [80]. Analogous to *B. mori*, we also identified 23 potential ILP paralogs in the nemertodermatid *Ascoparia* sp. (Figure 3). The nermertodermatids *N. westbladi* and *M. stichopi* possess three ILPs and one ILP, respectively. *X. profunda* has two ILPs, while *X. bocki* possesses only one. The only acoel species in which we identified a potential ILP is *D. gymnopharyngeus*, but this ILP shows an unusually 100 amino acid long C-chain that separates the two predicted active insulin chains, which is longer than the commonly 10-50 amino acids in the other xenacoelomorph and other metazoan ILPs (see Figure 3).

**Figure 3:**
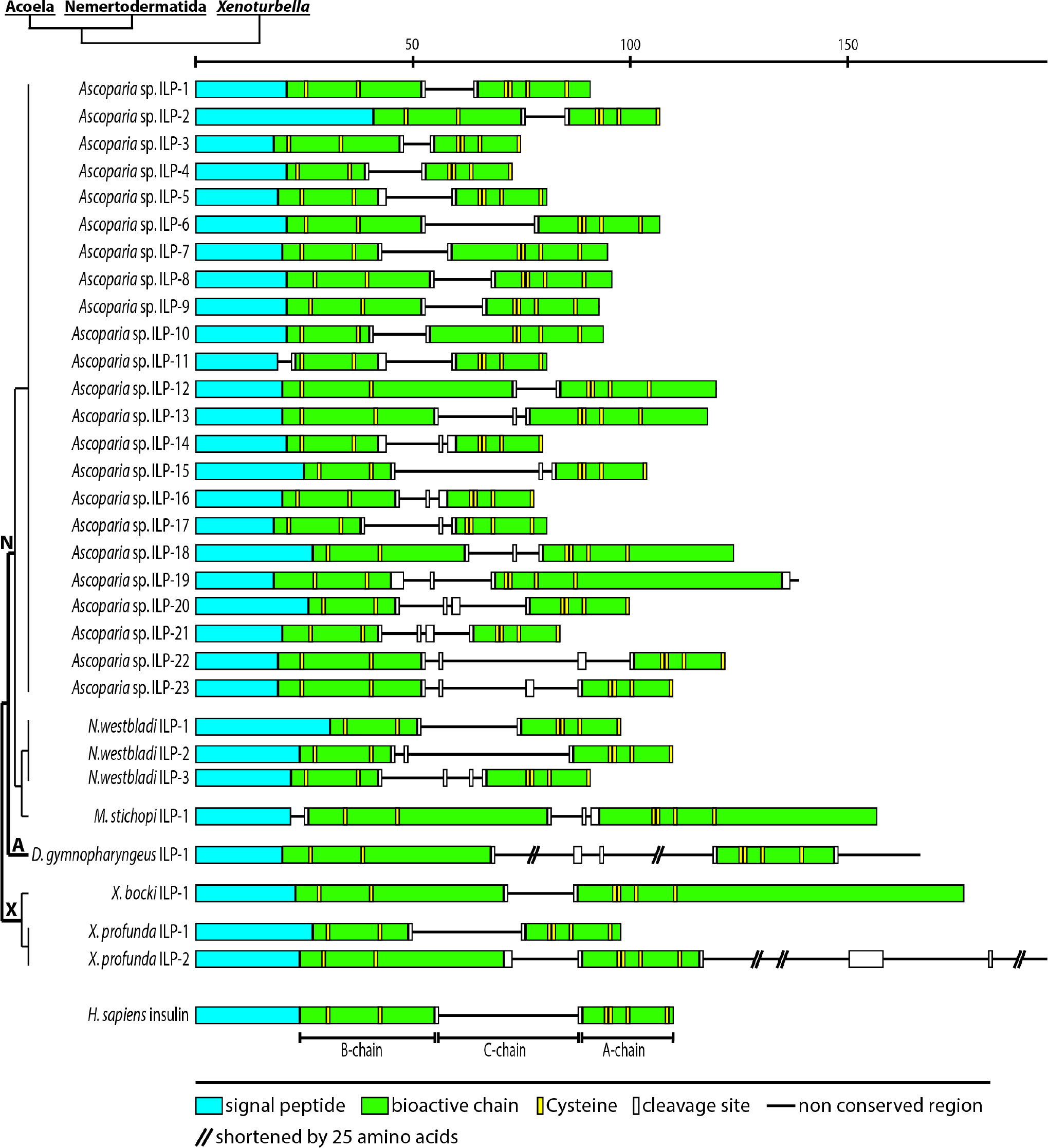
Xenacoelomorph insulin like peptides (ILPs). Schematic representation of precursor arrangement and comparison with human insulin. *H. sapiens* insulin precursor [P01308.1] consisting of B- and A-chain with conserved cysteine residues and a connecting C-chain. ILP insulin-like peptide, A Acoela, N Nemertodermatida, *X. Xenoturbella*. Scale bar above the precursor schemes indicates the length in number of amino acids.

#### Prokineticin related peptides

Prokineticin like peptides have been identified in nephrozoans and related peptides with similar cysteine arrangements have also been found in cnidarians [2]. We identified in both *Xenoturbella* species and in two nemertodermatids (i.e., *M. stichopi* and *N. westbladi*) a single prokineticin precursor (Supplementary Figure 1). However, we could not detect any prokineticin related peptide in any of the acoel species. While the receptors of glycoprotein hormones, insulins and the insulin related relaxin peptides are conserved, with clear orthologs in bilaterians and non-bilaterians, the prokineticin receptor is only known from deuterostomes, and no orthologs have been identified in protostomes or non-bilaterian animals [1, 2, 5, 81]. We identified an ortholog of the deuterostome prokineticin GPCR in the nemertodermatid *N. westbladi* (Figure 4, Supplementary Figures 2 and 3). This is the first evidence that the prokineticin GPCR has already been present in the last common ancestor of all bilaterians, which suggests its loss in protostomes.

**Figure 4:**
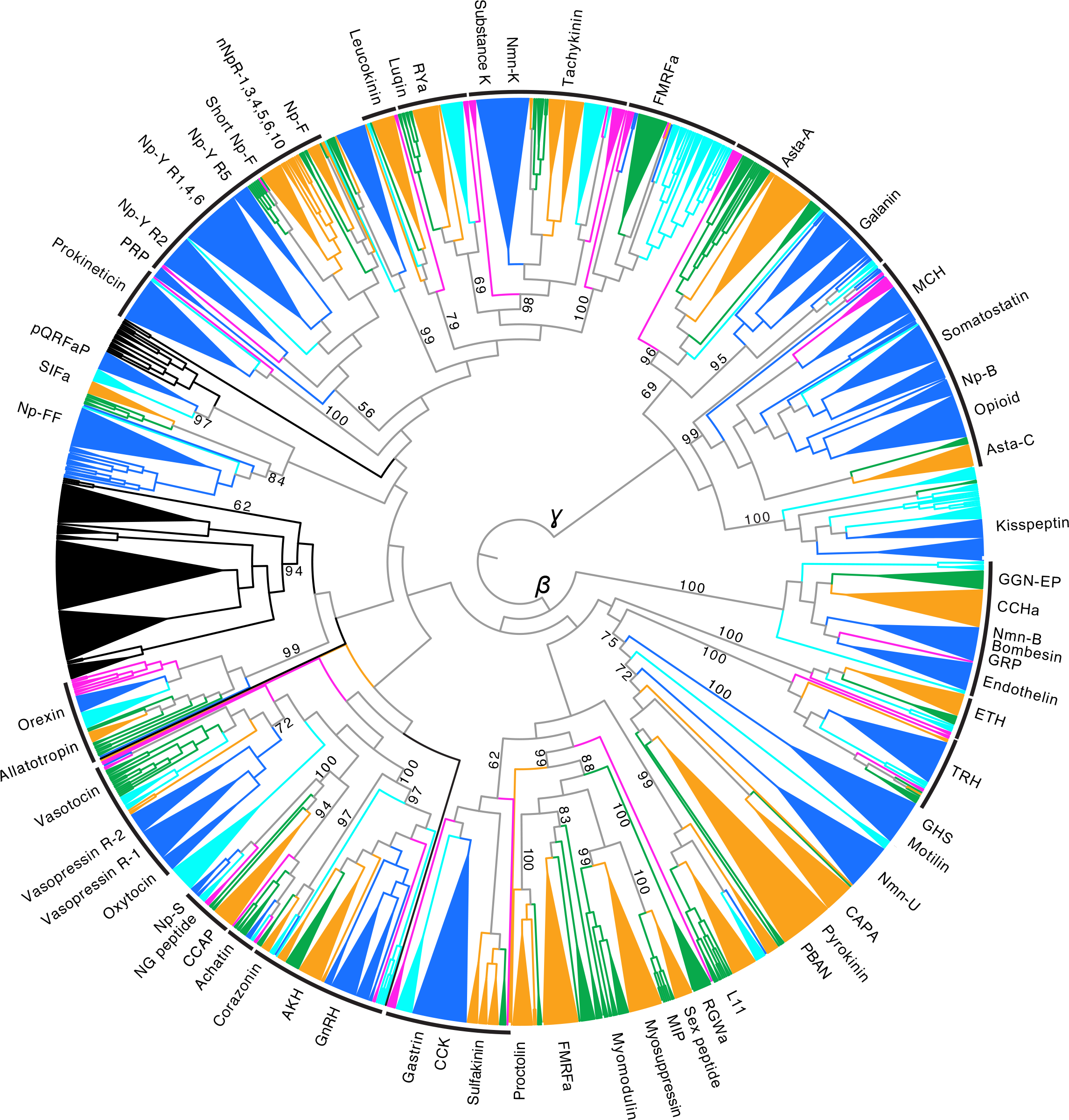
RAxML tree of Rhodopsin type neuropeptide GPCRs. Rhodopsin *beta* GPCRs are rooted against rhodopsin *gamma* GPCRs. Support values (bootstrap support with 100 bootstrap replicates) are shown for the major branches that constitute orthologues receptor types from different animal groups. Branches of homologues receptors that belong to the same animal group with support values of at least 80 are combined into a triangle. Black lines at the periphery of the tree indicate bilaterian receptor groups with at least one corresponding xenacoelomorph receptor. *a* amide, *AKH* adipokinetic hormone, *Asta* allatostatin, *CCAP* crustacean cardioaccelatory peptide, *CCK* cholecystokinin, *GGN-EP* GGN excitatory peptide, *ETH* ecdysis triggering hormone, *GHS* growth hormone segretagogue, *GnRH* gonadotropin releasing hormone, *GRP* gastrin releasing peptide, *Np* neuropeptide, *nNpR* nematode neuropeptide receptor, *Nmn* Neuromedin, *L11* elevenin, *MCH* melanin concentrating hormone, *MIP* myoinhibitory peptide, *PBAN* pheromone biosynthesis activating neuropeptide, *pQRFaP* pyroglutaminated RFamide peptide, *PRP* prolactin releasing peptide, *TRH* thryotropin releasing hormone. Colour coding: Magenta = xenacoelomorphs, dark blue = chordates, light blue = ambulacrarians, orange = ecdysozoans, green = spiralians, black = cnidarians.

### Xenacoelomorph orthologs of nephrozoan neuropeptide GPCRs reveal the ancient bilaterian signaling complement

In our GPCRs survey, we detected orthologs of most characterized nephrozoan neuropeptide GPCRs, revealing their presence in the last common ancestor of all bilaterians including Xenacoelomorpha (Figures 4 and 5, Supplementary Figure 2-5). The phylogenetic analysis of the GPCRs generally recovered the different groups of neuropeptide GPCRs that have previously been shown to be plesiomorphic to Deuterostomia and Protostomia [1, 2, 23]. The different types of rhodopsin- and secretin-type GPCRs (Figures 4 and 5) were confirmed in both phylogenetic analysis with trimmed sequences (Supplementary Figures 2 and 4) and in the cluster analysis with untrimmed sequences (Supplementary Figures 3 and 5). In total, we identified receptors that can be grouped into 20 ancestral bilaterian peptidergic systems, 19 of these receptors were present in nemertodermatids and 13 in *Xenoturbella* species, whereas only 8 conserved homologs were identified in Acoela (Figure 6).

**Figure 5:**
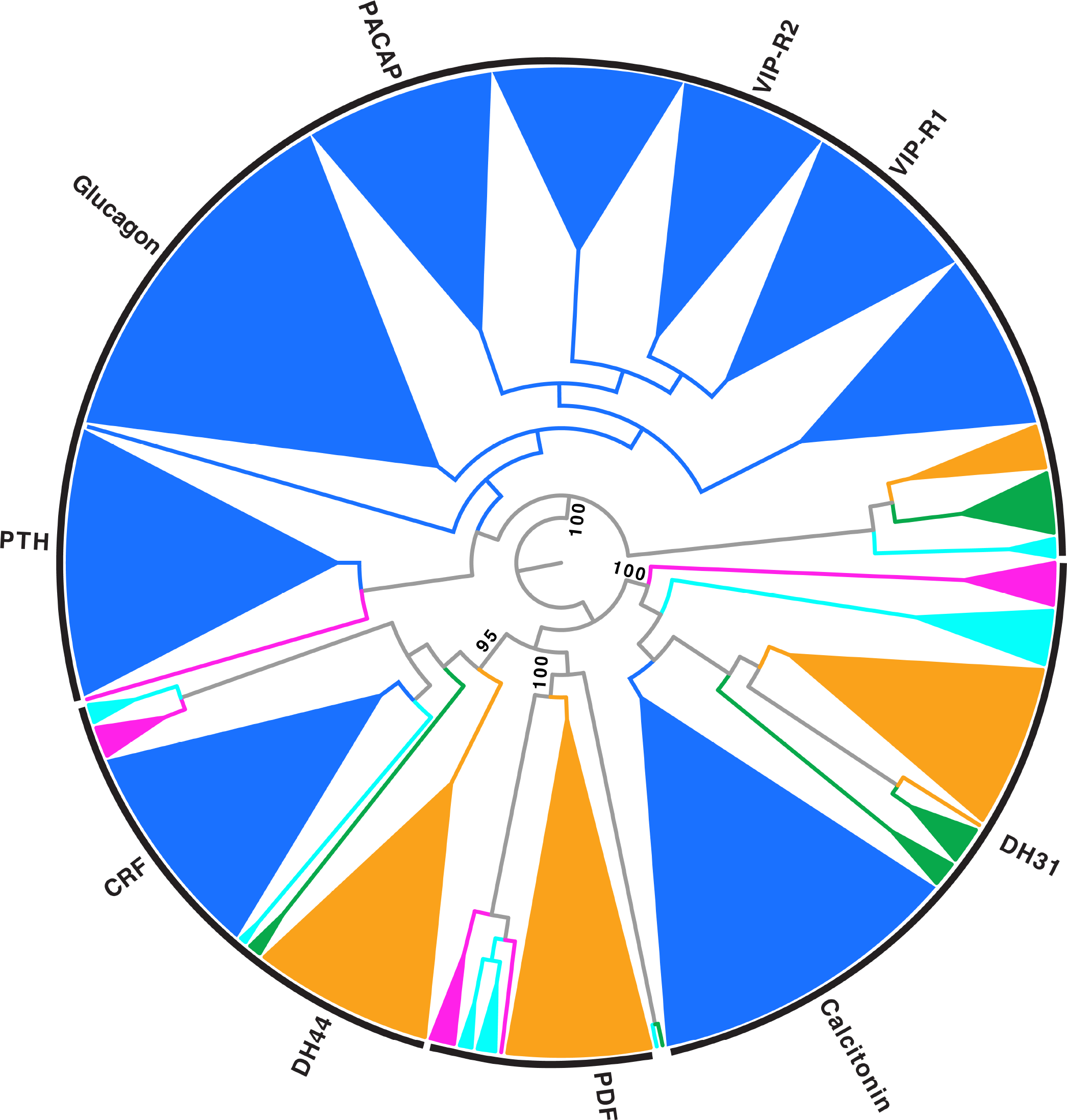
RAxML tree of secretin type neuropeptide GPCRs. Glucagon-related receptor group is rooted against the remaining receptors. Support values (bootstrap support with 100 bootstrap replicates) are shown for the major branches that constitute orthologues receptor types from different animal groups. Branches of homologues receptors that belong to the same animal grouping with support values of at least 80 are combined into a triangle. Black lines at the periphery of the tree indicate bilaterian receptor groups with at least one xenacoelomorph receptor. *CRF* corticotropin releasing factor, *DH* diuretic hormone, *PDF* pigment dispersing factor, *PTH* parathyroid hormone, *VIP* vasoactive intestinal peptide, *PACAP* pituitary adenylate cyclaseactivating polypeptide, -*R* receptor. Colour coding: Magenta = xenacoelomorphs, dark blue = chordates, light blue = ambulacrarians, orange = ecdysozoans, green = spiralians, black = cnidarians.

**Figure 6:**
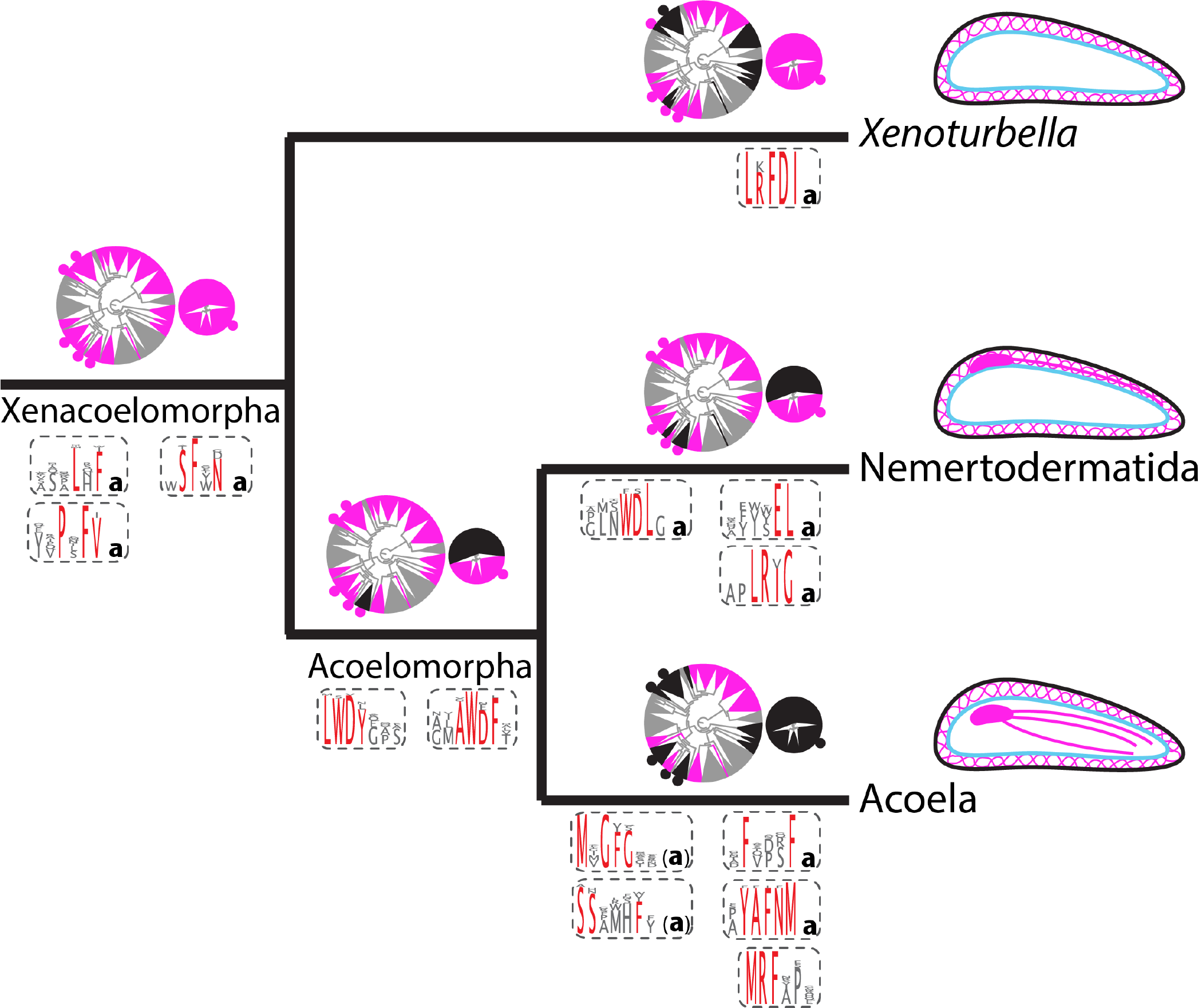
Lineage specific peptidergic systems of Xenacoelomorpha. Radial trees above branches show presence and absence of conserved bilaterian peptidergic rhodopsin type GPCRs (bigger tree on the left) and secretin type GPCRs (smaller tree on the right). Trees are derived from figure 4 and 5 with all branches of one receptor type collapsed into a single triangle. Conserved bilaterian ligands that were detected in Xenacoelomorpha are indicated by filled circles next to the corresponding receptor type (rhodopsin tree, top to bottom: neuropeptide Y/F, prokineticin, vasotocin, achatin, GnRH; secretin tree: calcitonin). Magenta indicates the presence of at least one corresponding sequence of this lineage and black indicates a lineage specific loss/absence. Peptide sequence logos below branches show lineage specific multi-copy peptides; logos were created by alignments of all paracopies from the different homologs. Conserved amino acids are highlighted red and represent peptide names, non-capitalized a at the end indicates amidation. Drawings on the right side (above lineage names) indicate simplified, lineage specific differences in nervous systems (magenta with its position in regards to epidermis (black and musculature (blue) in side view. (*Xenoturbella* with basiepidermal nerve net and without nerve condensation, Nemertodermatida with basiepidermal nerve net and basiepidermal brain and nerve cords, Acoela with basiepidermal nerve net and internalized brain and nerve cords.)

In addition to the previously described presence of the trochozoan FMRFamide-, tachykinin/neuromedin K- and luqin/RYamide-like receptors [53], we identified rhodopsin beta GPCRs of achatin, vasotocin/oxytocin/vasopressin, GnRH/AKH, orexin/allatotropin, leucokinin, gastrin/cholecystokinin/sulfakinin, neuropeptide S/NG peptide/crustacean cardio accelatory peptide, thyrotropin releasing hormone, ecdysis triggering hormone, GGN excitatory peptide/CCHamide/neuromedin B/endothelin receptors, and a single receptor that shows some affinity to a big group that diversified strongly in protostomes and includes proctolin, arthropod FMRFamide, myomodulin and myoinhibitory peptide receptors. We also found three orthologs of receptors from the cluster of the neuropeptide Y, neuropeptide F, short neuropeptide F and prolactin releasing peptide GPCRs, which grouped at two different positions. One sequence from *N. westbladi* and a sequence from *M. stichopi* seem to be closely-related to protostome short neuropeptide F and neuropeptide F receptors, whereas another sequence from *N. westbladi* clusters closer to the deuterostome neuropeptide Y receptor 2 group. These neuropeptide Y/F and related receptor groups show strong connections in the cluster analysis (Supplementary Figure 3) but the relationship of these groups was not well resolved in our analyses, similar to previous analyses [1, 2].

We also detected several rhodopsin gamma GPCRs, which cluster in the two groups of melanin concentrating hormone/opioid/somatostatin/allatostatin C receptors and the galanin/allatostatin A receptors. The first group seems to have expanded in deuterostomes but shows an unclear phylogeny and strong connections in the clustermap in our analysis as well as in previous analyses [1, 2]. The xenacoelomorph GPCRs of this group are devided into two distinct groups, one containing receptors from *N. westbladi* and *H. miamia* and the other one from *N. westbladi, H. miamia, M. stichopi, C. macropyga and D. gymnopharyngaeus*.

From the fewer groups of secretin-type GPCRs, we identified receptors that are related to Calcitonin/DH31, corticotropin releasing factor/diuretic hormone 44 and pigment dispersing factor signaling. We also detected a single sequence in *Xenoturbell bocki* that is related to the strongly diversified deuterostome glucagon/parathyroid hormone/vasoactive intestinal peptide receptors with a group of related protostome orphan GPCRs (Figure 5 and Supplementary Figures 4 and 5). In total, we found evidence for the presence of most receptors that are known to be shared between protostomes and deuterostomes, with only few exceptions. It has previously been shown that the last common ancestor of protostomes and deuterostomes likely already had two paralogs of the related gonadotropin releasing hormone (GnRH)/adipokinetic hormone (AKH) and corazonin systems [23]. We only identified one type of these systems in xenacoelomorphs, which showed more similarity to the GnRH/AKH receptors. These GPCRs however, were only discovered in the two *Xenoturbella* species and it cannot be ruled out that a corazonin-like paralog is part of the ancestral xenacoelomorph complement but was not detected in our survey, as a lack of detection in our transcriptomes might not necessarily mean a loss from the genomes. We also found a single type of GnRH/AKH/corazonin related ligand in nemertodermatids and *Xenoturbella* species (described below). We could not find any xenacoelomorph orthologs for neuromedin U/capability/pyrokinin, neuropeptide FF/SIFamide, elevenin and Kisspeptin signaling, which are otherwise present in protostomes and deuterostomes.

### Five conserved bilaterian preproneuropeptides are present in nemertodermatids and *Xenoturbella*, but absent in acoels

For five of the above listed neuropeptide GPCRs that are present in xenacoelomorphs, we identified precursors of potential ligands that show strong similarity to the corresponding nephrozoan neuropeptides: calcitonin, vasotocin-neurophysin, GnRH/AKH/corazonin, achatin and neuropeptide Y/F related precursors. We found these bilaterian neuropeptide precursors only in nemertodermatid and *Xenoturbella* species but not in any of the acoel transcriptomes. Their presence in xenacoelomorphs and their high similarity to protostomian and deuterostomian orthologs, shows that these ancient ligands have diverged less when compared to most other bilaterian neuropeptides.

#### *Calcitonin* (Figure 7)

Calcitonins are characterized by two conserved cysteine residues at the N-terminus of the active peptide and are usually encoded a single time on their precursors, with some distance between the signal peptide and the active peptide [1, 55]. We detected six calcitonin prepropeptides in nemertodermatids and one in *X. bocki*. While the nemertodermatids *N. westbladi* and *Ascoparia* sp. possess one calcitonin precursor each, the two more closely related species *M. stichopi* and *Sterreria* sp. have two paralogs that likely arose from an independent duplication event (Figure 7b).

**Figure 7:**
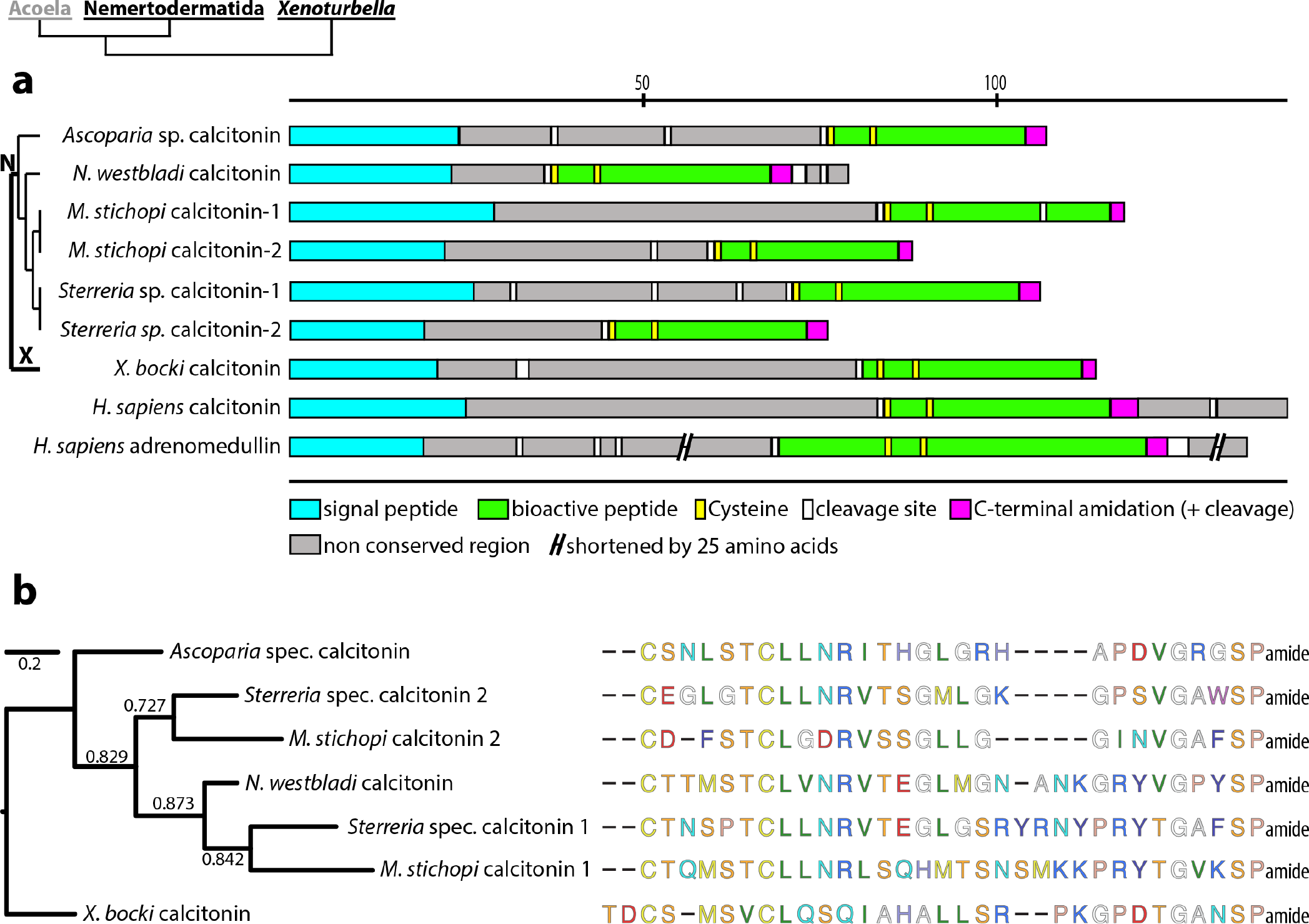
Xenacoelomorph calcitonin-like neuropeptides. **a)** Schematic representation of precursor arrangement, compared to human calcitonin [P01258.2] and human adrenomedullin [P35318.1]. **b)** FastTree phylogenetic analysis of predicted calcitonin-like ligands with nemertodermatid sequences rooted against the *X. bocki* sequence. Amino acids are coded according to their biochemical properties. Numbers on branches indicate SH-like local support. N Nemertodermatida, X *Xenoturbella*

#### *Vasotocin-neurophysin* (Figure 8)

Vasotocin-neurophysin related peptides have a conserved propeptide structure in which the shorter vasotocin peptide is adjacent to a longer neurophysin domain, both of which contain strongly conserved cysteine residues [1, 82]. We discovered three complete sequences of vasotocin-neurophysin precursors: one sequence in the nemertodermatid *Ascoparia* sp. and one in each *Xenoturbella* species. An alignment with 39 oxytocin-/vasopressin-/vasotocin-neurophysin sequences from different bilaterian taxa shows that a high conservation of the cysteine residues in both domains (Figure 8b) is also present in Xenacoelomorpha.

**Figure 8:**
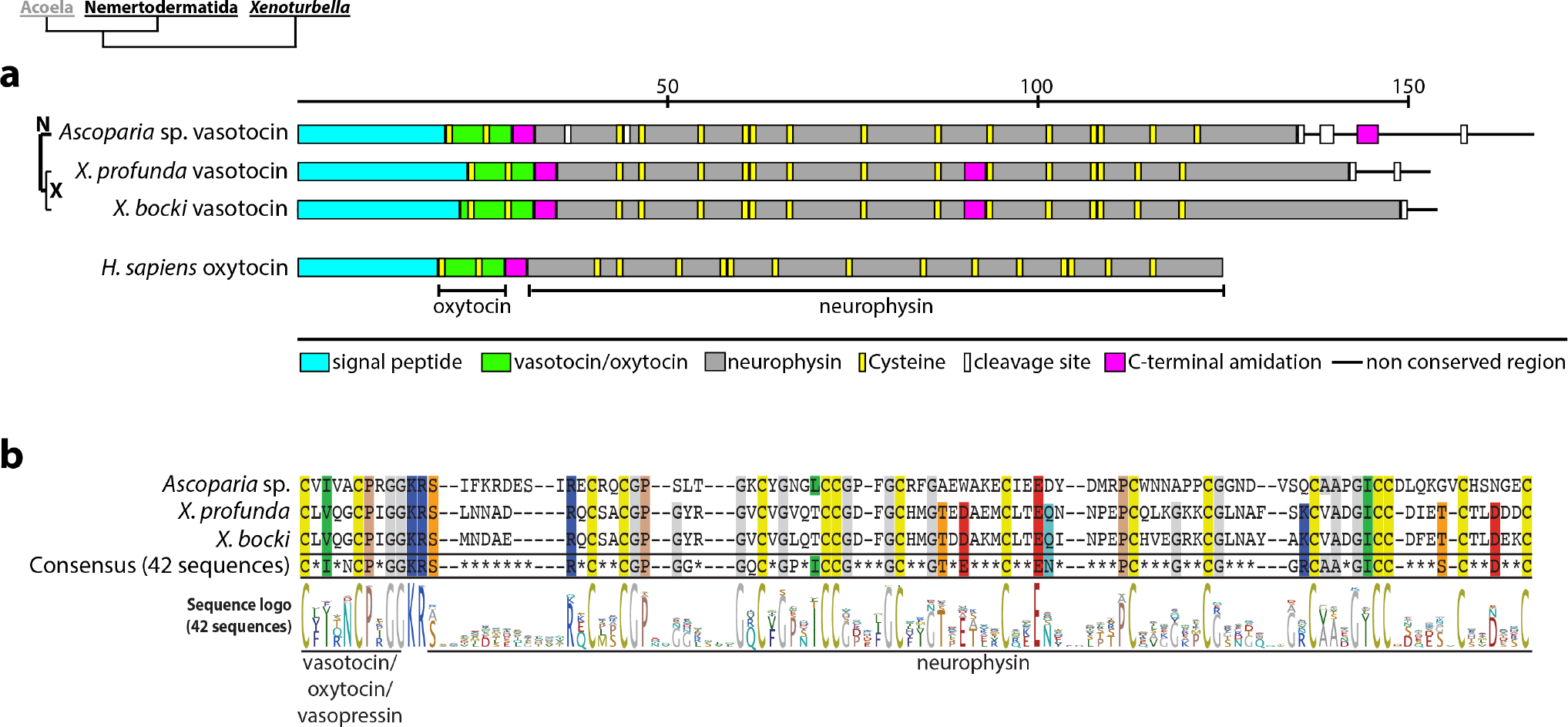
Xenacoelomorph vasotocin. **a)** Schematic precursor arrangement of xenacoelomorph vasotocins, compared to human oxytocin. **b)** Alignment with consensus sequence and peptide sequence logo created from 42 sequences of different bilaterian vasotocin-/oxytocin-/vasopressin-related peptides, including the three xenacoelomorph sequences. *H. sapiens* oxytocin [Accession P01178.1]. N Nemertodermatida, X *Xenoturbella*

#### *GnRH-/AKH-/corazonin-like peptide* (Figure 9)

It has previously been shown that gonadotropin releasing hormone (GnRH), adipokinetic hormone (AKH) and corazonin (CRZ) signaling are homologues to each other [83-86]. The overall sequence similarity between these related peptides is not very high but the prepropeptide arrangements are identical and the active ligands usually have their N-terminus capped by N-pyroglutamyl formation and their C-terminus by amidation [22, 83, 87-89]. Exceptions are the nematode AKH-related peptides that lack the C-terminal amidation [85, 87] or the non-pyroglutaminated ligand that activates the *Asterias rubens* CRZ receptor homolog [23]. We discovered neuropeptide precursor candidates in several nemertodermatid and both *Xenoturbella* species that showed similarity to GnRH, AKH and CRZ peptides. The predicted peptide starts after the signal peptide with a length of 13 amino acids (15 amino acids in one of the *N. westbladi* paralogs), followed by a non-conserved region with two cysteine residues. All predicted peptides are amidated, however, an N-terminal glutamine is not present. One of the *N. westbladi* paralogs underwent an evolutionary modification and differs from other bilaterian sequences in having two GnRH/AKH/CRZ-related peptides after the signal peptide.

**Figure 9:**
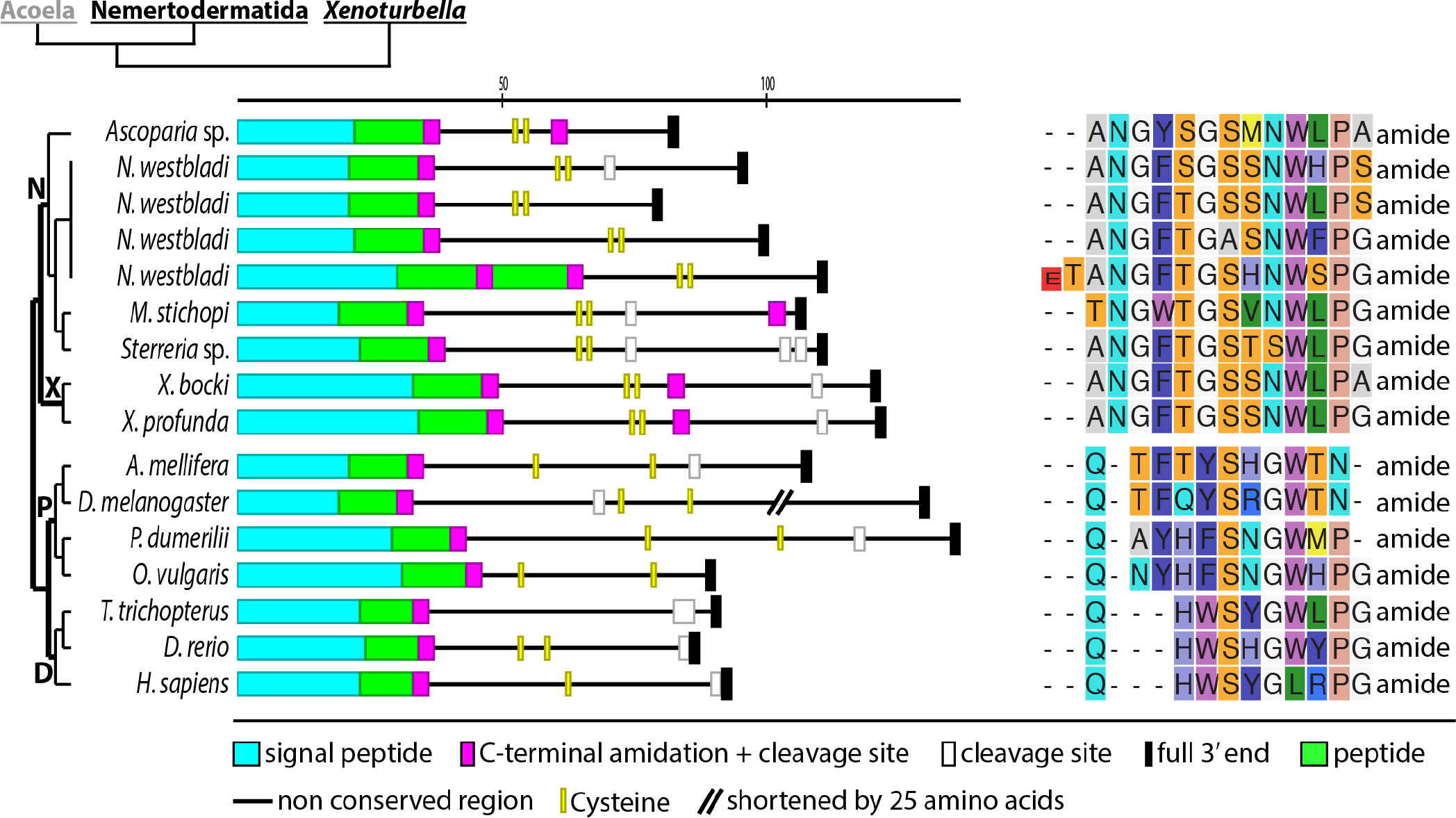
Xenacoelomorph GnRH- /corazonin-related neuropeptides. Schematic representation of peptide precursor sequences and alignment of predicted active peptides with mixed corazonin and GnRH-like ligands of deuterostomes and protostomes. D Deuterostomia, N Nemertodermatida, P Protostomia, *X. Xenoturbella*.

#### *Achatin* (Figure 10)

Achatins can be found in different non-vertebrate deuterostomes and in many protostomes, and they seem to represent an exceptionally well conserved type of short multi-copy peptide, which consists of four amino acids with an aromatic D-amino acid in the second position [2, 59, 90]. In the transcriptomes of both *Xenoturbella* species and in those of the nemertodermatids *M. stichopi* and *N. westbladi*, we identified precursors for the achatin-like peptide GFGN (Figure 10). The precursors of both *Xenoturbella* species are similar, with five repeats of GFGN on the C-terminus, separated from the signal peptide by nearly 60 amino acids. The achatin-like peptides of *M. stichopi* and *N. westbladi* are more dispersed in their precursors. The sequence of the predicted achatin peptides is identical to achatin peptides of the ambulacrarian *Saccoglossus kowalevskii*, the cephalochordate *Branchiostoma floridae* as well as to the achatins of the ecdysozoans *Priapulus caudatus* and *Halicryptus spinulosus* (see Figure 10).

**Figure 10:**
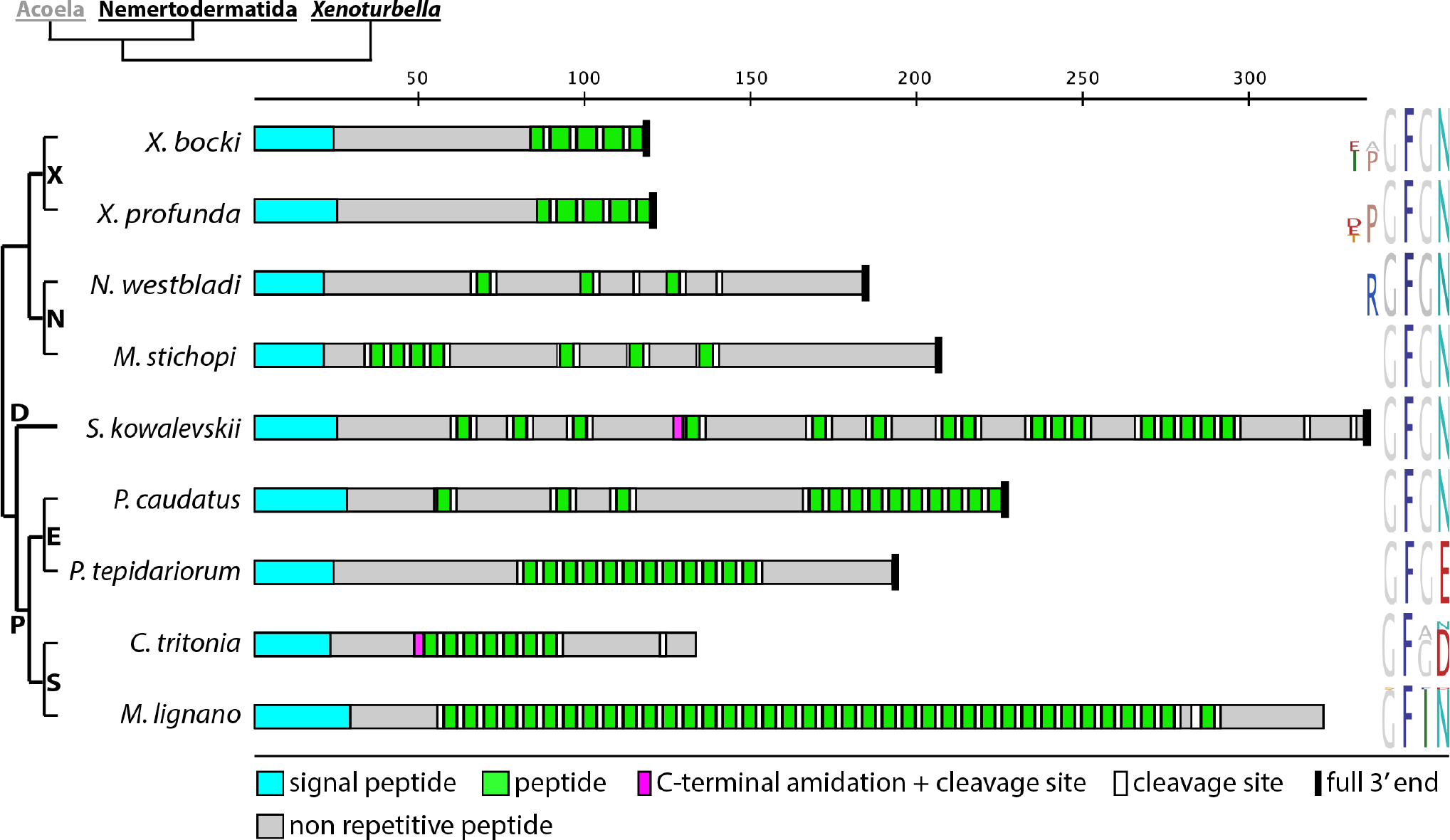
Schematic representation of achatin precursors with peptide sequence logos of the predicted peptides. Peptide sequence logos are based on the alignment of all indicated peptide paracopies from each precursor. Reference sequences are *Saccoglossus kowalevskii* (XP_002732147.1), *Priapulus caudatus* (XP_014662331.1), *Parasteatoda tepidariorum* (XP_015908358.1), *Charonia tritonis* (AQS80481.1) and *Macrostomum lignano* (PAA66935.1). D Deuterostomia, E Ecdysozoa, N Nemertodermatida, P Protostomia, S Spiralia, X *Xenoturbella*.

#### *Neuropeptide Y/F* (Figure 11)

Neuropeptide Y is known from deuterostomes and is the ortholog of the protostome neuropeptide F, both of which are medium long peptides that are encoded a single time on their precursor [1, 2, 91]. The only potential homologs of neuropeptide Y or neuropeptide F has been identified in the transcriptome of *N. westbladi*. Two of the three candidates were fully recovered. They all share the C-terminal amino acid residue arginine-proline-arginine, followed by one of the two aromatic amino acids tyrosine or phenylalanine. Whether the ancestral Xenacoelomorph neuropeptide Y/F ended in a phenylalanine or a tyrosine cannot be reconstructed. Interestingly, the C-terminal YYAIVGRPRFamide motif of one of the *N. westbladi* homologs is similar to the C-terminus found in various protostomes such as certain platyhelminth- and trochozoan-species, indicating that a very similar motif might have been present in the ancestral bilaterian neuropeptide Y/F [55, 57, 91-93]. The predicted peptide length of two of the *N. westbladi* paralogs is shorter than what is usually predicted for many nephrozoan neuropeptide Y/Fs. However, there are also some bilaterian exceptions such as the *Apis melifera* and *Lottia gigantea* neuropeptide F, which have two basic amino acids at a similar position, and the neuropeptide F of *C. gigas*, where an alternative monobasic cleavage site has even been confirmed by mass spectrometry [91, 92, 94].

**Figure 11:**
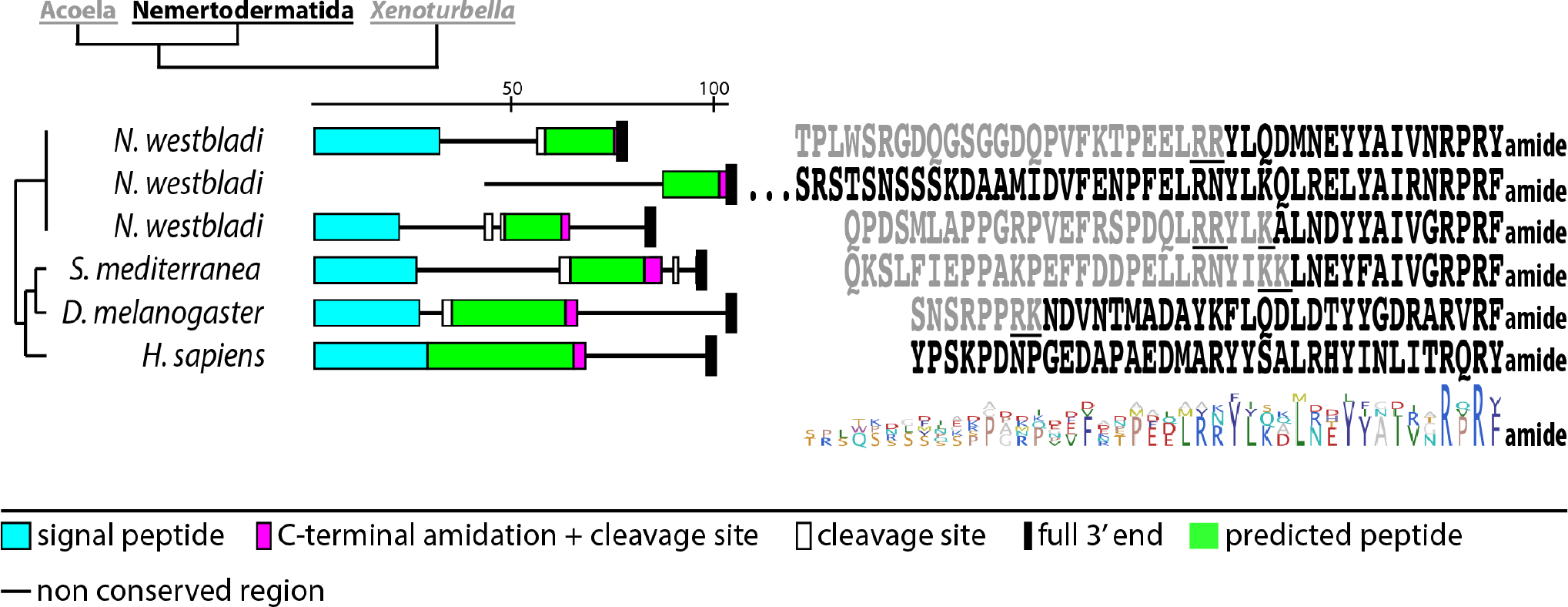
Nemertodermatid neuropeptide Y/F. Schematic representation of precursor with alignment of the predicted peptides. Black amino acids represent active peptides, underlined amino acids indicate predicted cleavage sites. *Homo sapiens* [P01303.1], *Schmidtea mediterranea* [ADC84429.1], *Drosophila melanogaster* [Q9VET0.1].

### Novel xenacoelomorph MCPs and their clade specific distribution

Beside the neuropeptides that are known from other animals, we identified a variety of multi-copy peptides (MCPs) that seem to be specific to all Xenacoelomorpha or certain xenacoelomorph groups. Only very few of them showed some similarity to clade-specific protostome or deuterostome multi-copy peptides. We also confirmed the presence of many of the peptide groups in acoels by mass spectrometry. We identified three types of peptides that are plesiomorphic for Xenacoelomorpha and which are present in *Xenoturbella* and Acoelomorpha species: LxFamides, SFxNamides and PxFVamides – with ‘x’ being a non-conserved amino acid that varied between the peptides of different species. Two types of peptides are plesiomorphic for Acoelomorpha that show a conserved AWDF and LWDY motif. Five types of peptides were only shared between different Acoela species, sharing the motifs SSxxxF(amide), MxGF, MRF, FxxxFamide or FNMamide. In the nemertodermatid species, we identified three types of peptides, sharing the motifs LRIGamide, ELamide or WDLamide and only a single type of peptide was specific for the two *Xenoturbella* species, which shared the motif LRFDIamide (see Figure 1 and 6 for a summary).

### Xenacoelomorph MCPs

#### *SFxNamide peptides* (Figure 12)

Several peptide precursors that were identified in representatives of all three xenacoelomorph clades share the common sequence SFxNamide (‘x’ indicates a variable amino acid). The precursors have 3-8 repeats of a 4-5 amino acid peptide with a high conservation within each precursor. The propeptide arrangement of both *Xenoturbella* species is highly similar with partially conserved residues in the non-repetitive peptides between the SFWNamide repeats, but two additional copies in *X. bocki*. Overall, the repeats are distributed unevenly in the propeptides and are either separate from each other or in small clusters of two or three repeats. An exception is the propeptide of *Ascoparia* sp. where all four repeats are encoded directly next to each other at the end of the propeptide sequence.

**Figure 12:**
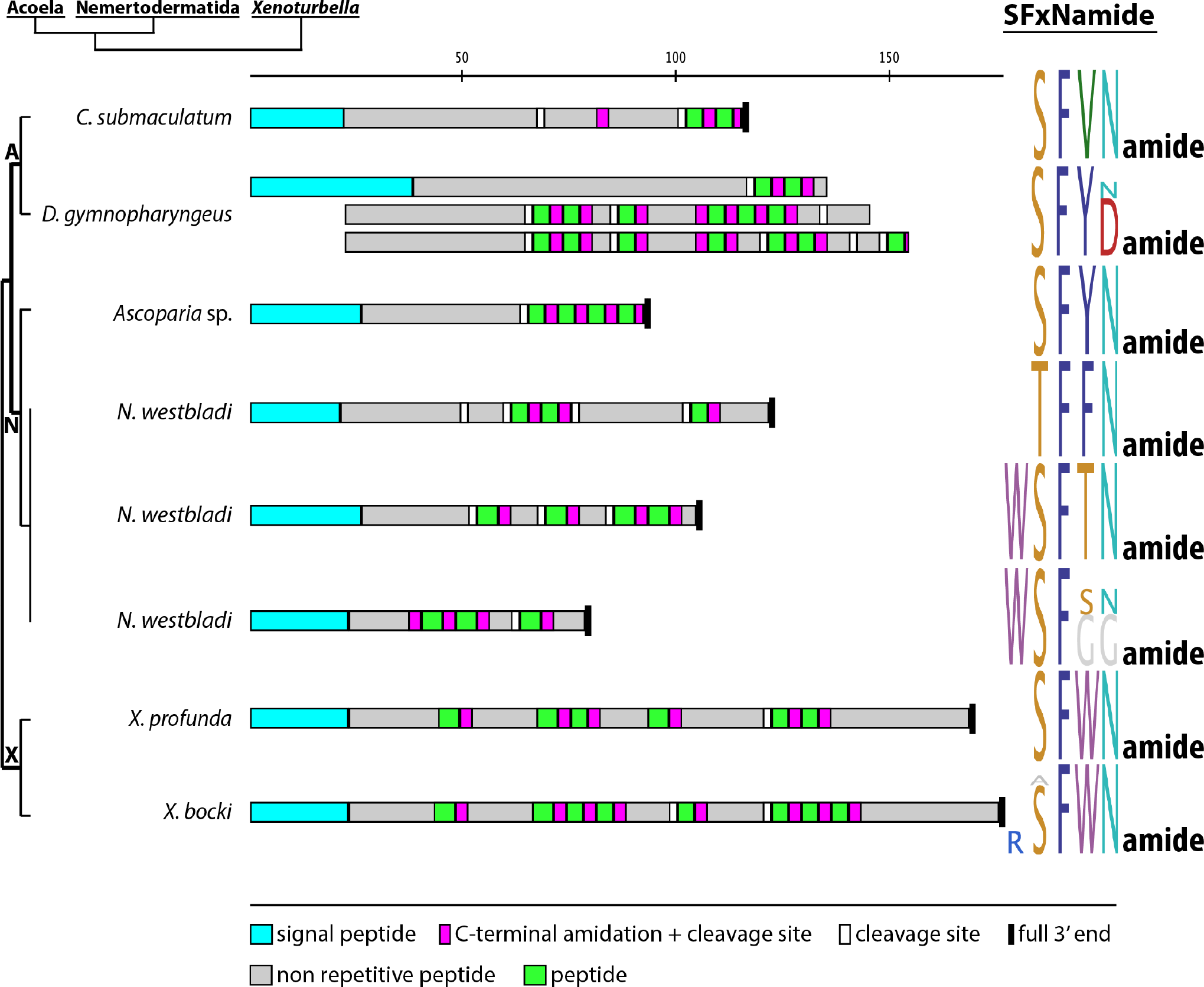
Xenacoelomorph SFxNamide peptides. Precursor scheme and peptide sequence logos of aligned peptides of each precursor. A Acoela, N Nemertodermatida, X *Xenoturbella*. Scale bar above precursor schemes indicates length in amino acids.

#### *LxFamide peptides* (Figure 13)

We detected in transcriptomes of Acoela, Nemertodermatida and *Xenoturbella* neuropeptides that share the C-terminal ending LxFamide. The peptide precursors of the fully recovered sequences show 4-5 repeats in different arrangements. The repeats are arranged in a cluster at the end of the propeptide in some of the propeptides (e.g., *I. pulchra* QPLNFamide and *D. gymnopharyngeus* LxFamides) and are separated by non-conserved amino acid sequences of different lengths in other propeptides (e.g., *D. longitubus* and *H. miamia*). The *D. gymnopharyngeus* ASALHFamide and *D. longitubus* ASAL[S/H]Famide are very similar in their repetitive sequences, but the precursor arrangement differ strongly. In *D. longitubus*, every repeat is separated by a non-repetitive part, whereas in *D. gymnopharyngeus*, the repeats are in one cluster. The sequence of *Ascoparia* sp. differs from the other fully recovered sequences in being nearly twice as long with 281 amino acids. Several LxFamides show similarity to echinoderm L-type SALMFamide (Figure 14).

**Figure 13:**
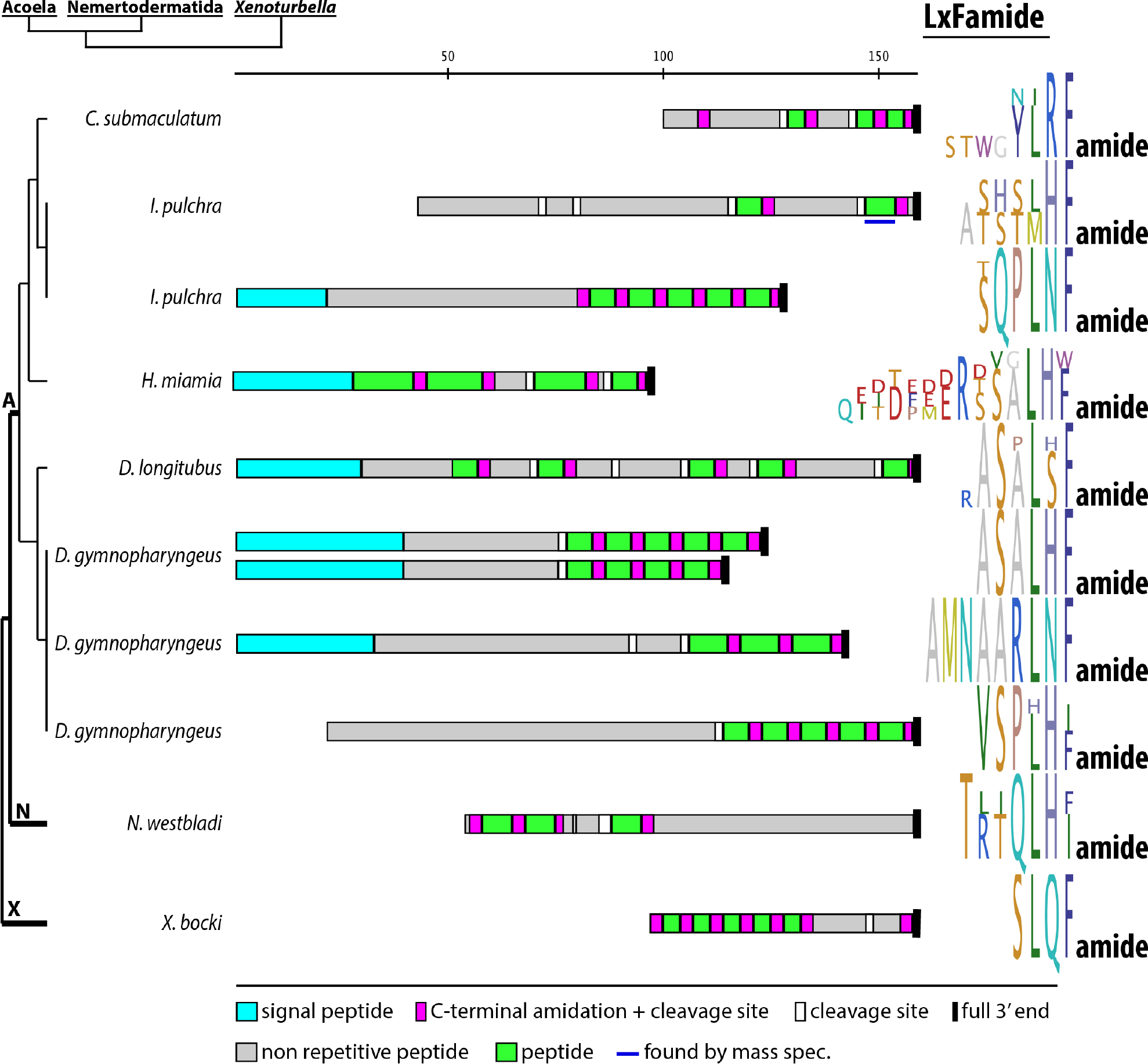
Xenacoelomorph LxFamide peptides. Precursor scheme and peptide sequence logos of aligned peptides of each precursor. A Acoela, N Nemertodermatida, X *Xenoturbella*. Scale bar above precursor schemes indicates length in amino acids.

**Figure 14:**
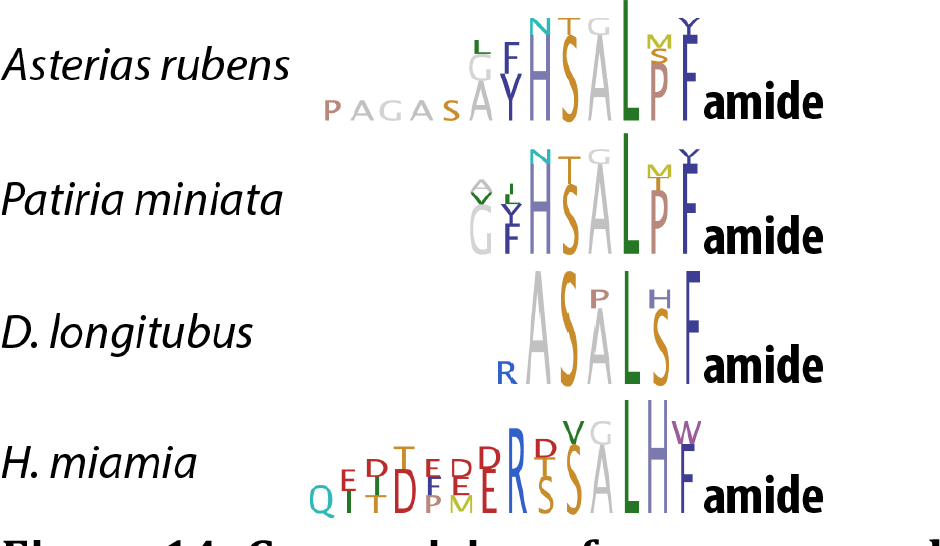
Comparision of some xenacoelomorph LxFamides and echinoderm SALMFamides. Peptide sequence logos were created from the alignment of the predicted neuropeptides. *Asterias rubens* L-type SALMFamide (ALJ99974.1) and *Patiria miniata* L-type SALMFamide published by Elphick et al 2013.

#### *PxFVamide peptides* (Figure 15)

We discovered two precursors in *N. westbladi* and one precursor in *X. bocki*, which encode short peptides with the common motif PxFVamide. Due to the lack of a 5’ end in one of the *N. westbladi* precursors and the 3’ end in the other one, the two *N. westbladi* fragments might belong to a single precursor. The peptide sequence logos are highly similar except for one additional amino acid residue in the 5’ end-missing transcript. The sequence discovered in *X. bocki* has only two repeats of the peptide and is clearly shorter than each of the two *N. westbladi* fragments. No sequence with the PxFVamide motif has been discovered in any of the acoel species. Multi-copy peptides with the C-terminal motif PxFVamide are also known from several molluscs (Figure 16).

**Figure 15:**
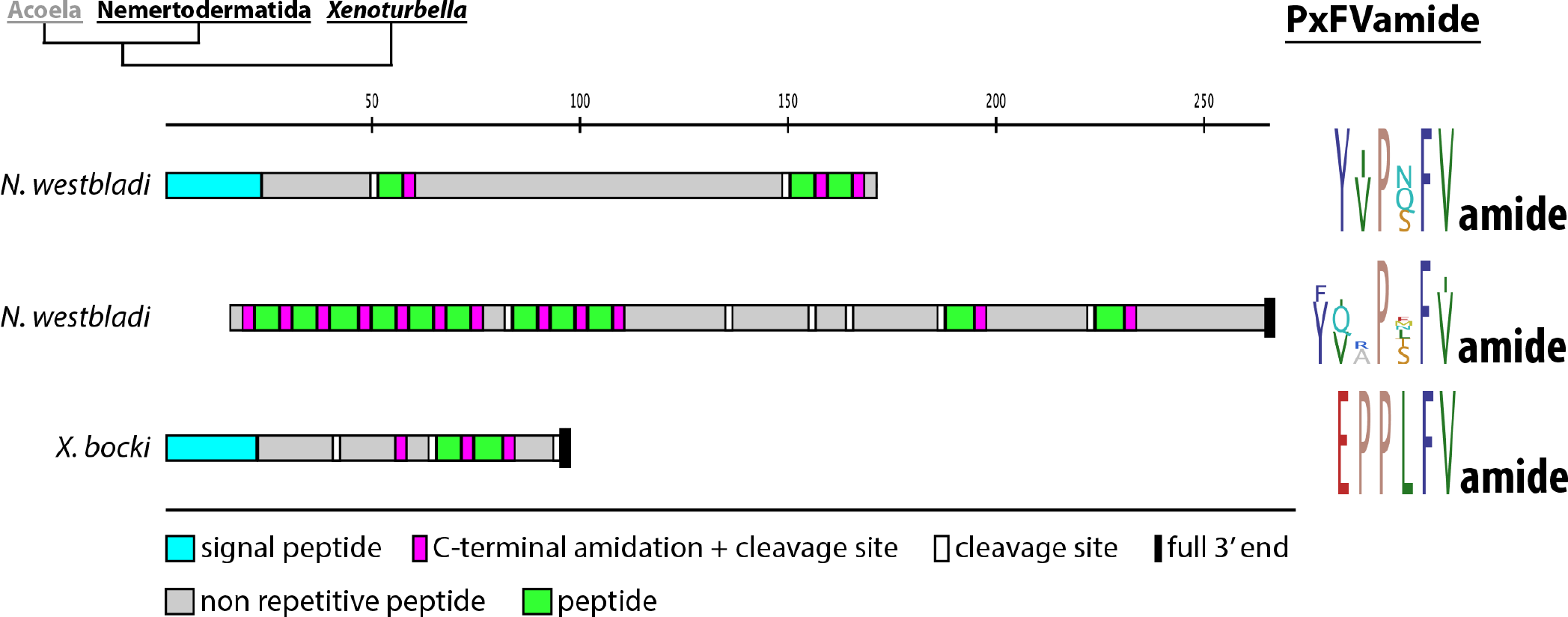
Xenacoelomorph PxFVamide peptides. Precursor scheme and peptide sequence logos of aligned peptides of each precursor. Scale bar above precursor schemes indicates length in amino acids.

**Figure 16:**
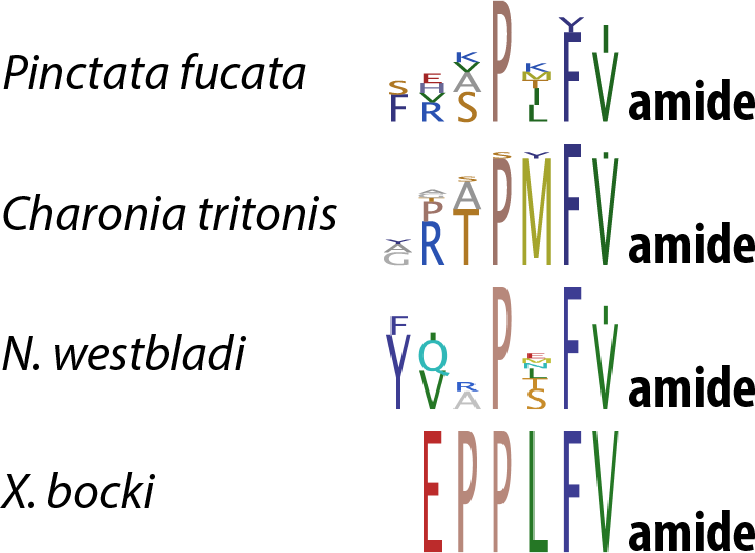
Comparision of xenacoelomorph PxFVamide peptides and some mollusc PxFVamide peptides. Peptide sequence logos were created from the alignment of the predicted neuropeptides. *Pinctata fucata* Mytilus inhibitory peptide, published by Steward et al. 2014, Charonia tritonis PXFVamide published by Bose et al. 2017 [AQS80535.1].

### Acoelomorph specific MCPs

#### *AWD and LWD peptides* (Figure 17)

We discovered two groups of peptides, AWD and LWD peptides that share the residues [tryptophan]-[aspartic acid]-[tyrosine/phenylalanine] and are both present in acoels and nemertodermatids. The sequence we detected in *D. longitubus* differs from the others in having a leucine after the aspartic acid instead of an aromatic amino acid. The AWDF peptides share a similar N-terminal extension [G/A]-[M/L]-A whereas the LWDY peptides have an extended C-terminus without obvious sequence conservation. Most peptide precursors have 3-9 repeats of the repeated peptides, which are separated from each other by a single, non-repetitive peptide. The LWD propeptide of *N. westbladi* differs in its arrangement from the other pecursors by having many non-seperated LWD repeats in one cluster. The AWDF peptide precursors of *C. submaculatum* and *N. westbladi* both have their first repeat with the additional C-terminal residues GGNamide, while the other repeats are shorter and without amidation sites. We could confirm peptides from the AWD as well as LWD propeptides that were flanked by basic cleavage sites by mass spectrometry. The similarities between the AWD and LWD sequences might indicate a duplication of an ancestral WD[F/Y] peptide, followed by a diversification into an AWD-like and LWD-like peptide, which would have happened before the split into Acoela and Nemertodermatida.

**Figure 17:**
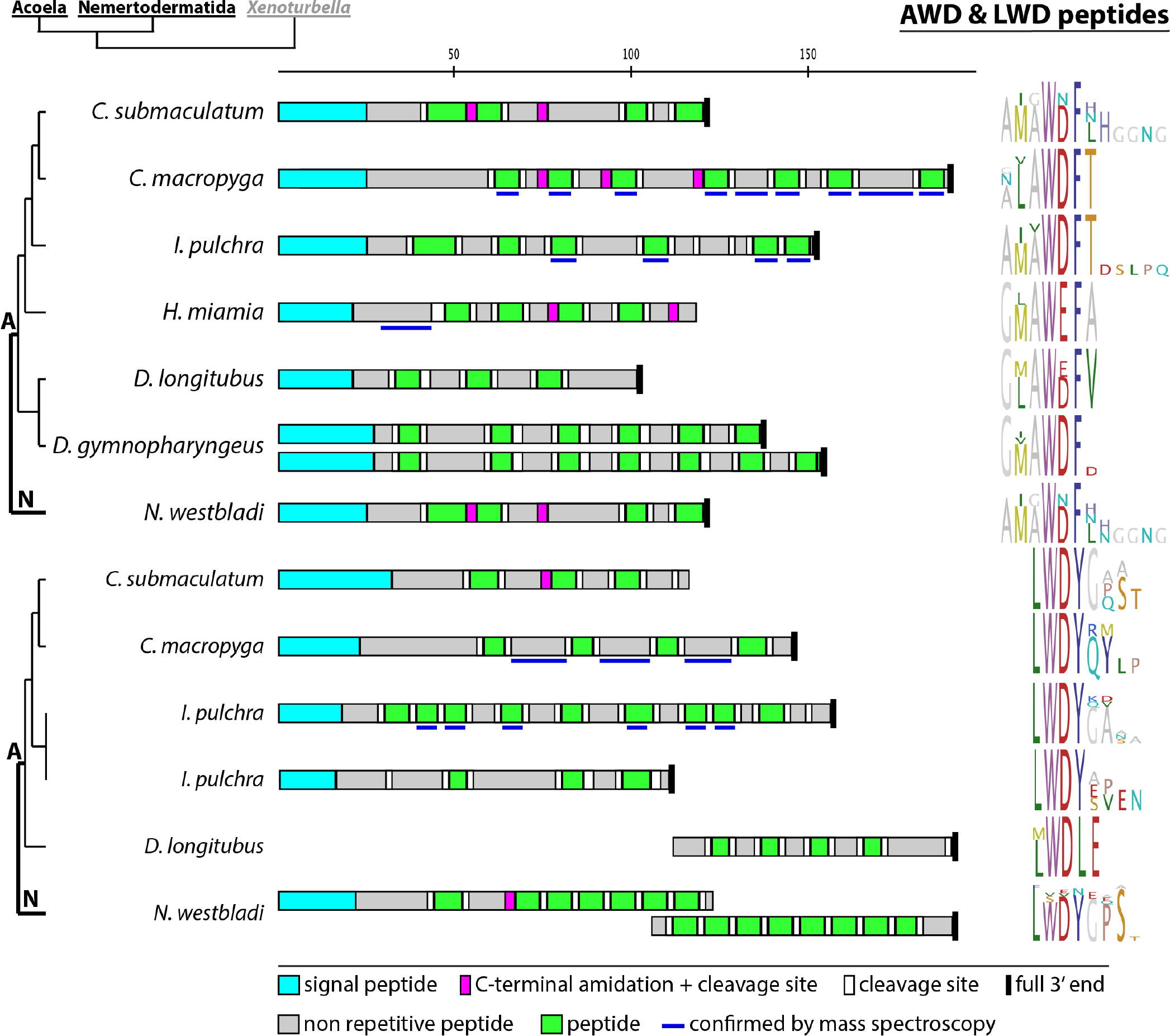
Acoelomorph AWD and LWD peptides. Precursor scheme and peptide sequence logos of aligned peptides of each precursor. A Acoela, N Nemertodermatida. Scale bar above precursor schemes indicates length in amino acids.

### Acoel specific MCPs

#### *SSxxxF peptides* (Figure 18)

Prepropeptides that share the repetitive sequence SSxxxF were discovered in the acoels *I. pulchra*, *E. macrobursarium* and *C. macropyga*. One of the three *C. macropyga* precursors and the precursors of *I. pulchra* and *E. macrobursalium* have nearly identical non-amidated SSAMHF[F/Y] peptides, but the *E. macrobursalium* and the *C. macropyga* peptides have additional, structurally related but amidated peptides. A mixture of structurally related amidated and non-amidated paracopies on the same precursor is rather rare and most multi-copy peptides encode either amidated or non-amidated paracopies. The amidated peptides are in one cluster in *C. macropyga*, whereas the non-amidated and amidated peptides in *E. macrobursalium* are alternating and they are mostly separated by less conserved sequences, which shows the general variability of the precursor structure of related orthologues peptides. The SSxxxF peptides show strong structural similarity to the LxFamides of *D. longitubus, D. gymnopharyngeus* and *H. miamia*, but do not share the leucine residue of the LxFamides, except for one of the amidated *E. macrobursalium* paracopies, indicating a homologues relationship between LxFamide and SSxxxF peptides, with several paralogs in *I. pulchra*, some of which show the SSxxxFamide motif, and others that show the LxFamide motif.

**Figure 18:**
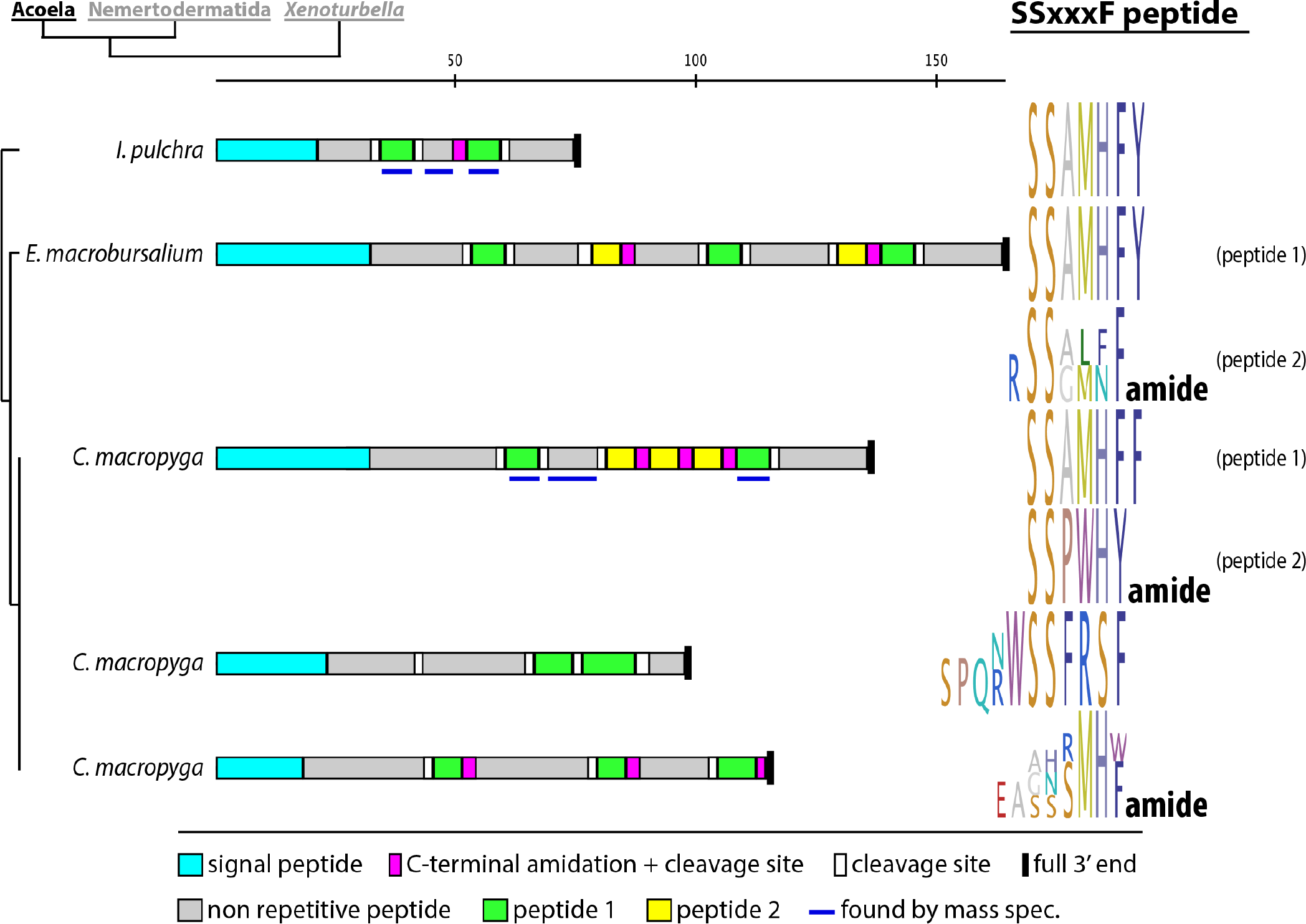
Acoel SSxxxF(amide) peptides. Precursor scheme and peptide sequence logos of aligned peptides of each precursor. Scale bar above precursor schemes indicates length in amino acids.

#### *MxGFG peptides* (Figure 19)

We detected precursor sequences with peptides that share the N-terminal motif MxGFG in several acoel species. In *C. submaculatum*, the aromatic phenylalanine residue at position 4 is exchanged by the aromatic tyrosine. Two different 3’ ends were found in *E. macrobursalium*, both of which contain several repeats of two different versions of the MxGFG, with one being amidated on the C-terminus but not the other. All repeated peptides in the precursors of *C. submaculatum* and *I. pulchra* show amidation sites, but both precursors are missing their 3’ end, whereas the two precursors with missing 5’ end in *E. macrobursalium* encode amidated as well as non-amidated repeats of similar peptides. The four transcripts found in *H. miamia, D. longitubus* and *D. gymnopharyngeus* show no amidation sites. Several predicted peptides from MxGFG prepropeptides were detected by mass spectrometry in *H. miamia* and *I. pulchra*. The MxGFG peptides share the GFG motif with the Nemertodermatid and *Xenoturbella* GFGN peptides and might be the acoel orthologs of achatin.

**Figure 19:**
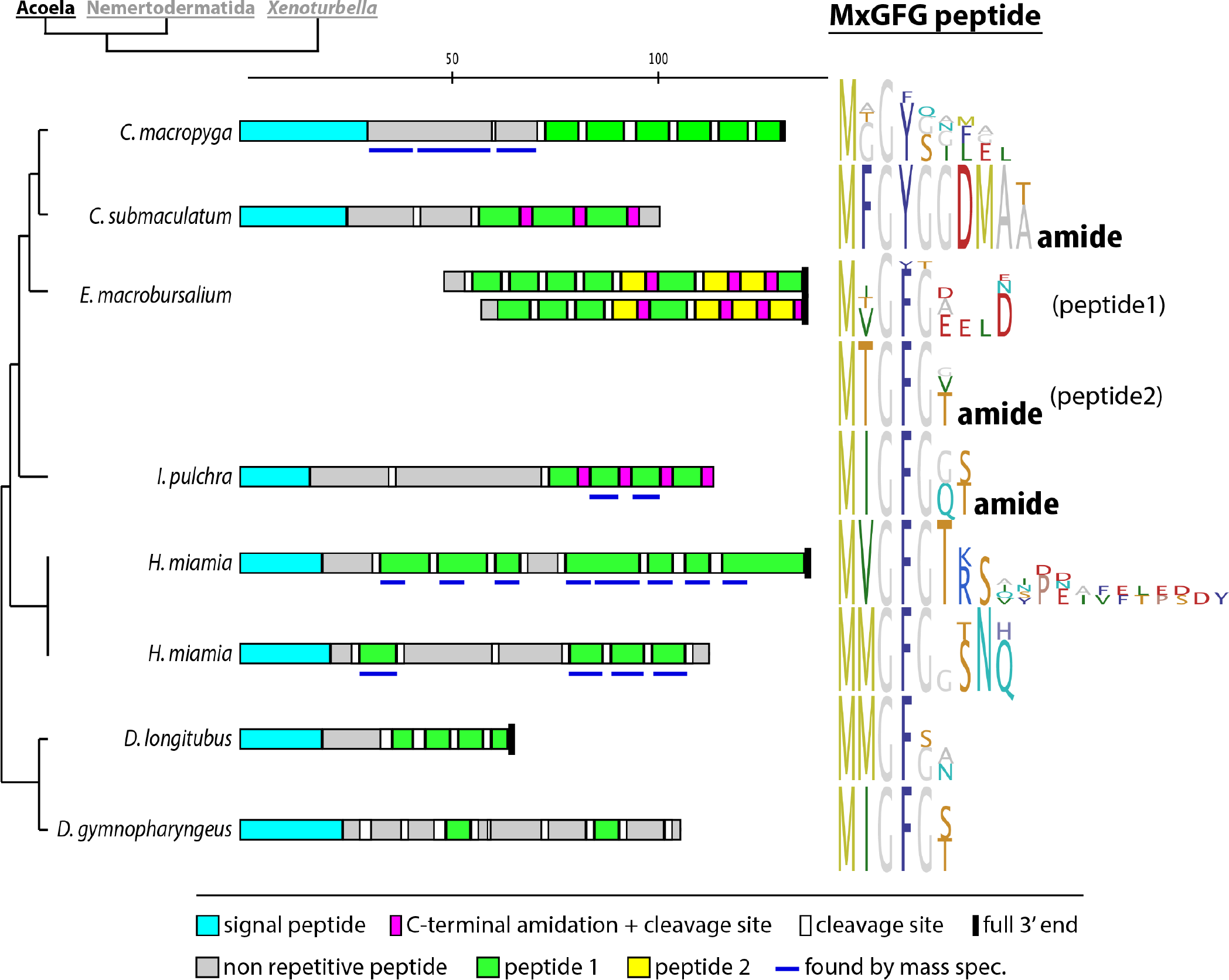
Acoel MxGFG peptides. Precursor scheme and peptide sequence logos of aligned peptides of each precursor. Scale bar above precursor schemes indicates length in amino acids.

#### *MRF peptides* (Figure 20)

In *I. pulchra* and *C. submaculatum*, we discovered precursor sequences that encode 6-7 peptides starting with the N-terminal sequence MRFxP. The peptides directly follow each other in both precursor sequences, being separated from the signal peptide by a short non-repetitive peptide in *I. pulchra* and three non-repetitive peptides in *C. submaculatum*. The C-terminal part of the peptides is neither conserved between the two species, nor within the same precursor.

**Figure 20:**
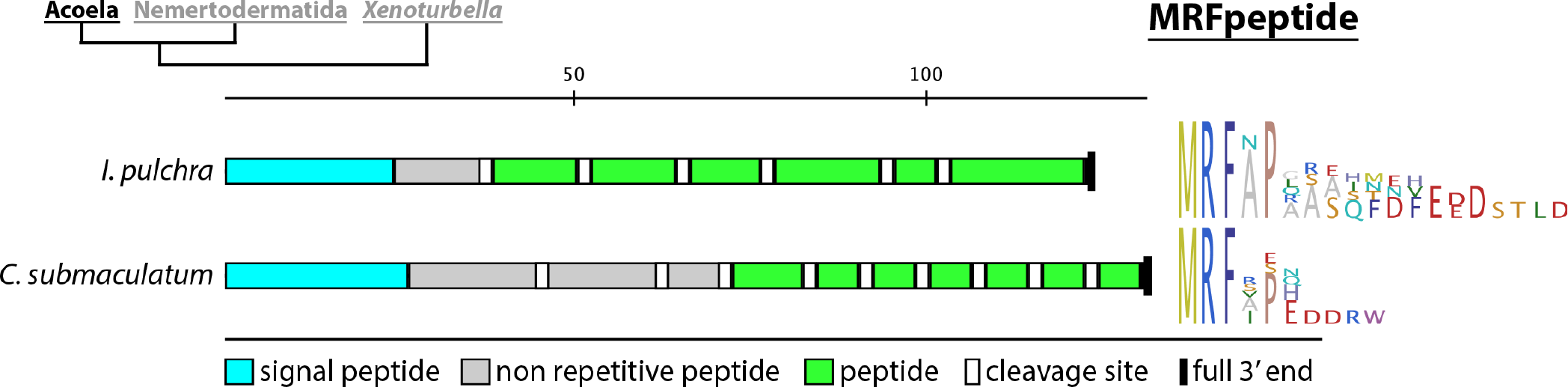
Acoel MRF peptides. Precursor scheme and peptide sequence logos of aligned peptides of each precursor. Scale bar above precursor schemes indicates length in amino acids.

#### *FxxxFamide peptides* (Figure 21)

In five acoel species, we found precursors that encode repetitive peptides with the conserved residues FxxxFamide. The three peptides of the closely-related *C. macropyga, C. submaculatum* and *I. pulchra* - at least in some of their copies - share the common sequence FxPSFamide, while the precursors of *D. longitubus* and *D. gymnopharyngeus* share the common motif FVDxFamide. The precursor schemes do not share any obvious similarities between each other except that the repetitive peptides are encoded in two clusters on the precursors of *C. macropyga* and *C. submaculatum*.

**Figure 21:**
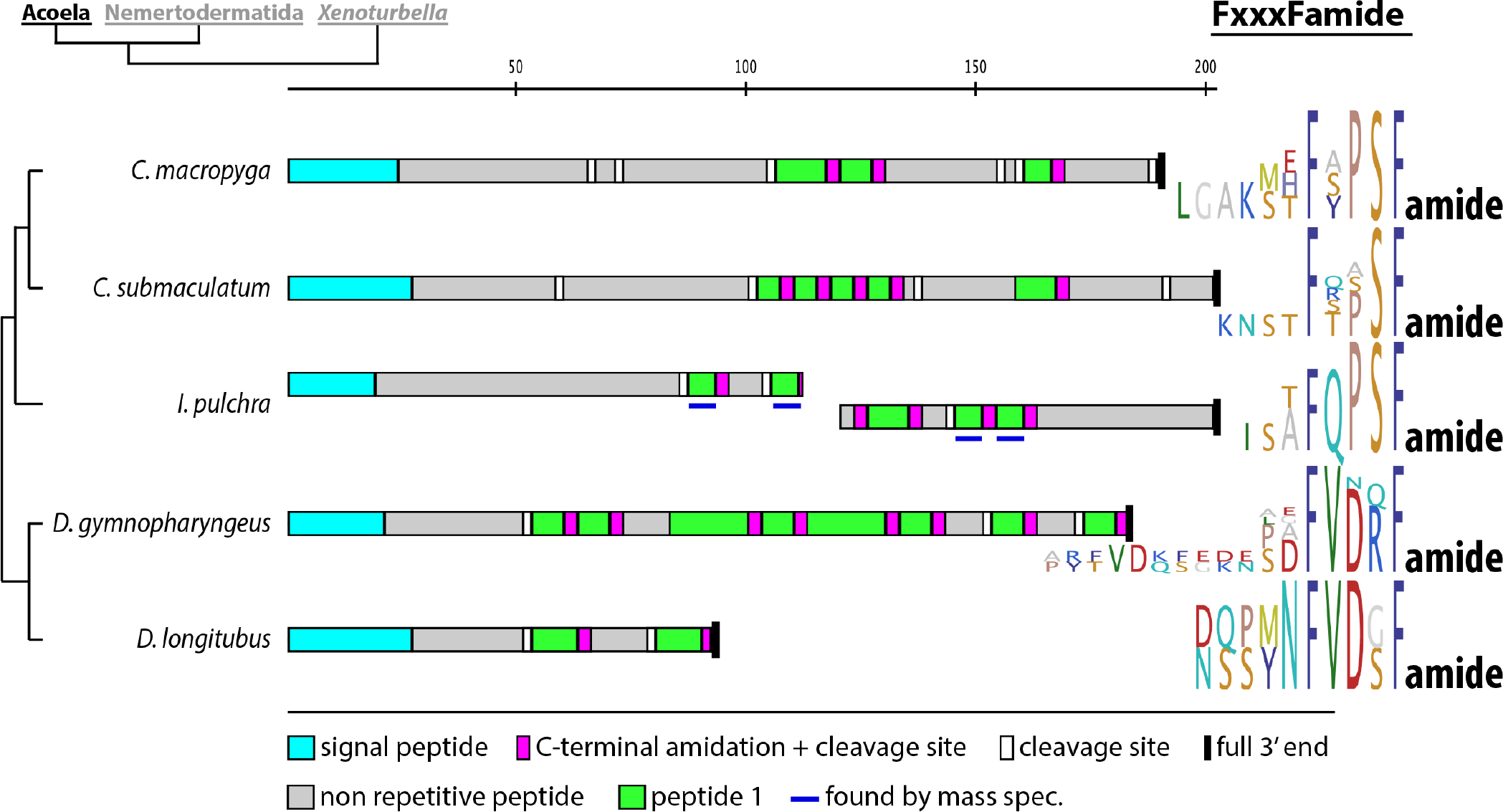
Acoel FxxxFamide peptides. Precursor scheme and peptide sequence logos of aligned peptides of each precursor. Scale bar above precursor schemes indicates length in amino acids.

#### *FNMamide peptides* (Figure 22)

We found precursor sequences encoding the peptide YAFNMamide in the acoel transcriptomes of *C. submaculatum, D. gymnopharyngeus* and *D. longitubus*. Except for the first few amino acids of the signal peptide, the precursor sequences of *C. submaculatum* and *D. gymnopharyngeus* are identical. The different FNMamide paracopies of the *C. submaculatum* and *D. gymnopharyngeus* precursors are nearly identical to each other.

**Figure 22:**
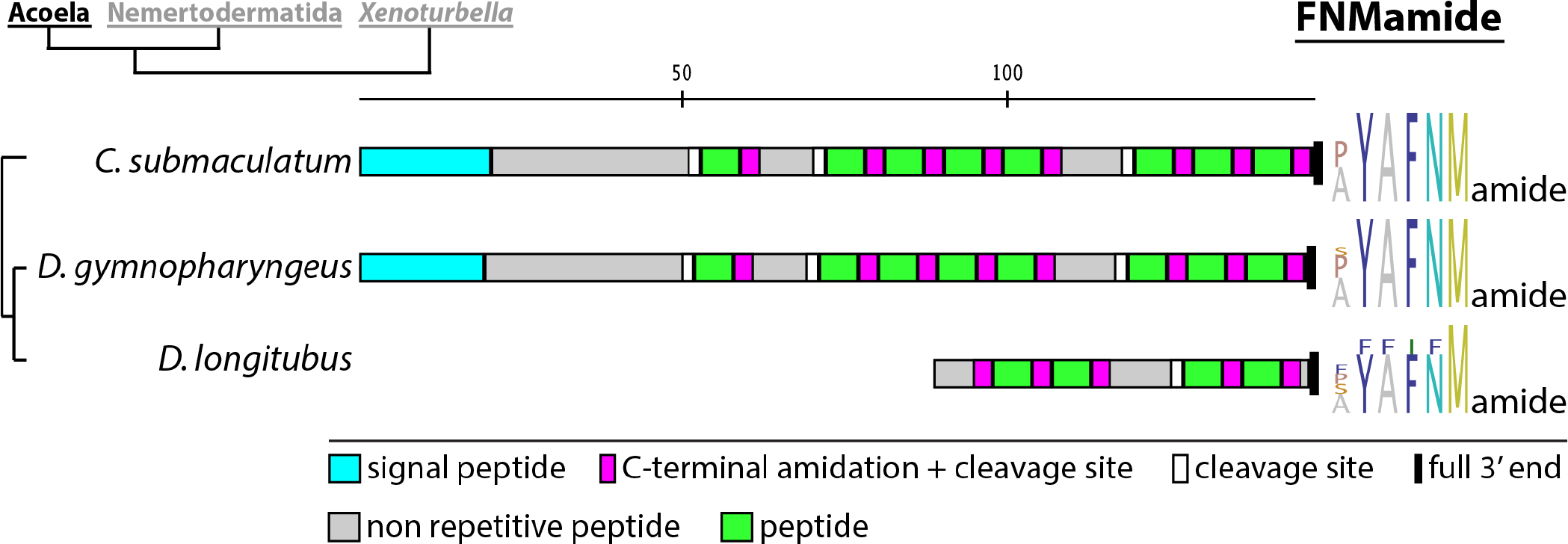
Acoel FNMamide peptides. Precursor scheme and peptide sequence logos of aligned peptides of each precursor. Scale bar above precursor schemes indicates length in amino acids.

### Nemertodermatid specific MCPs

#### *WDLamide/WDLGamide peptides* (Figure 23)

We discovered in all four nemertodermatid transcriptomes sequences with WDLamide/WDLGamide repeats. The sequences that we found in *Ascoparia* sp., *M. stichopi* and *Sterreria* sp. end with an amidated leucine, while the sequences of *N. westbladi* have an additional, amidated glycine residue on their C-terminus. The *N. westbladi* WDLGamide peptides share otherwise more sequence similarity to the *Ascoparia* sp. WDLamide than the WDLamide peptides of *Ascoparia* sp., *M. stichopi* and *Sterreria* sp. to each other.

**Figure 23:**
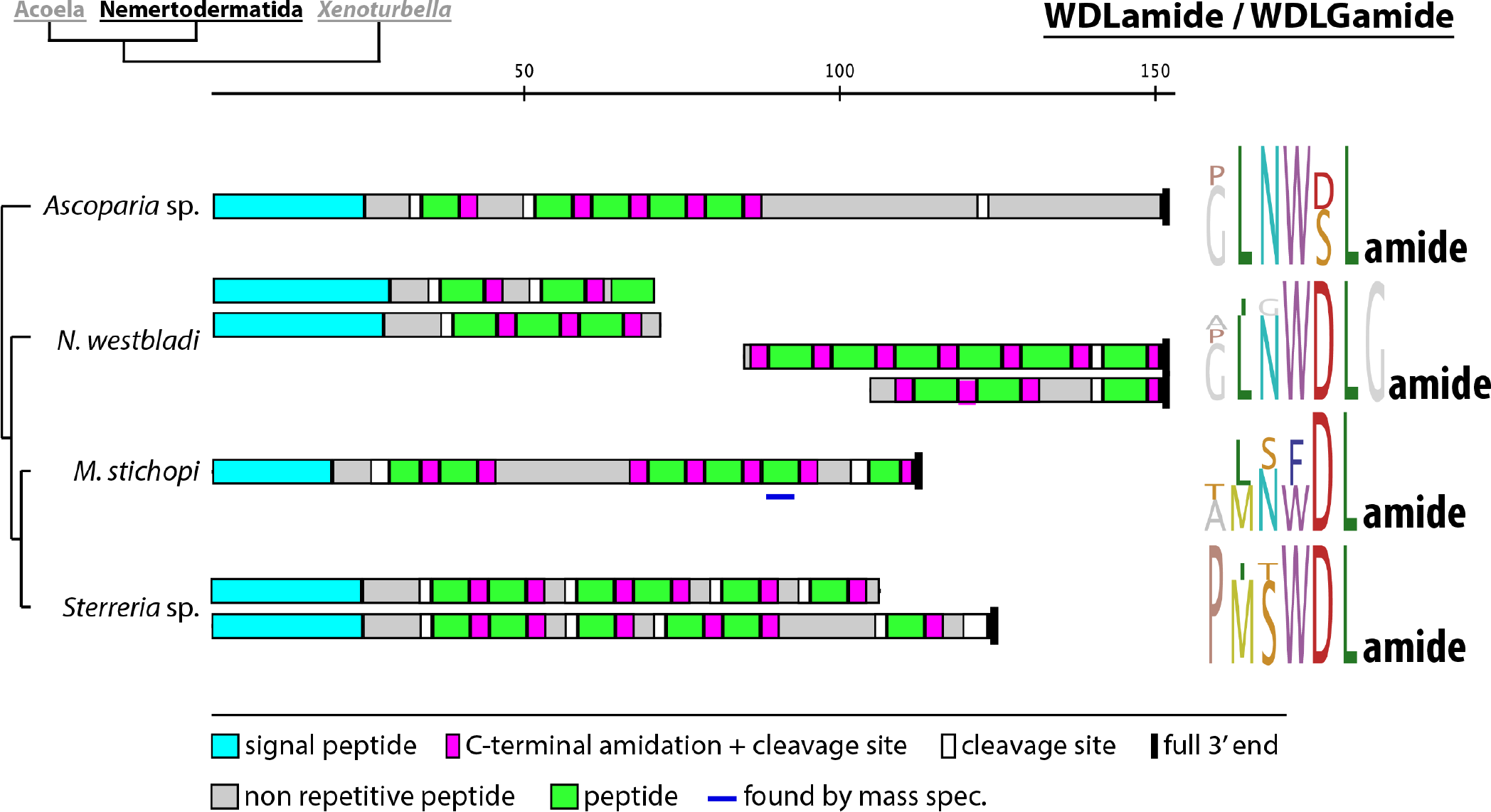
Nemertodermatid WDLamide/WDLGamide peptides. Precursor scheme and peptide sequence logos of aligned peptides of each precursor. Scale bar above precursor schemes indicates length in amino acids.

#### *ELamide peptides* (Figure 24)

In three nemertodermatid transciptomes, we found precursors that encode repetitive parts ending in ELamide. *N. westbladi* has two different paralogs, both with three repetitive peptides separated from each other by longer stretches of non-repetitive peptides, and the first repeat starting right after the signal peptide. The propeptide arrangement of *Sterreria* sp. and *Ascoparia* sp. ELamides are different from the *N. westbladi* ones in having several consecutive copies. The ELamide peptides show a certain similarity to arthropod allatostatin A, as further discussed later.

**Figure 24:**
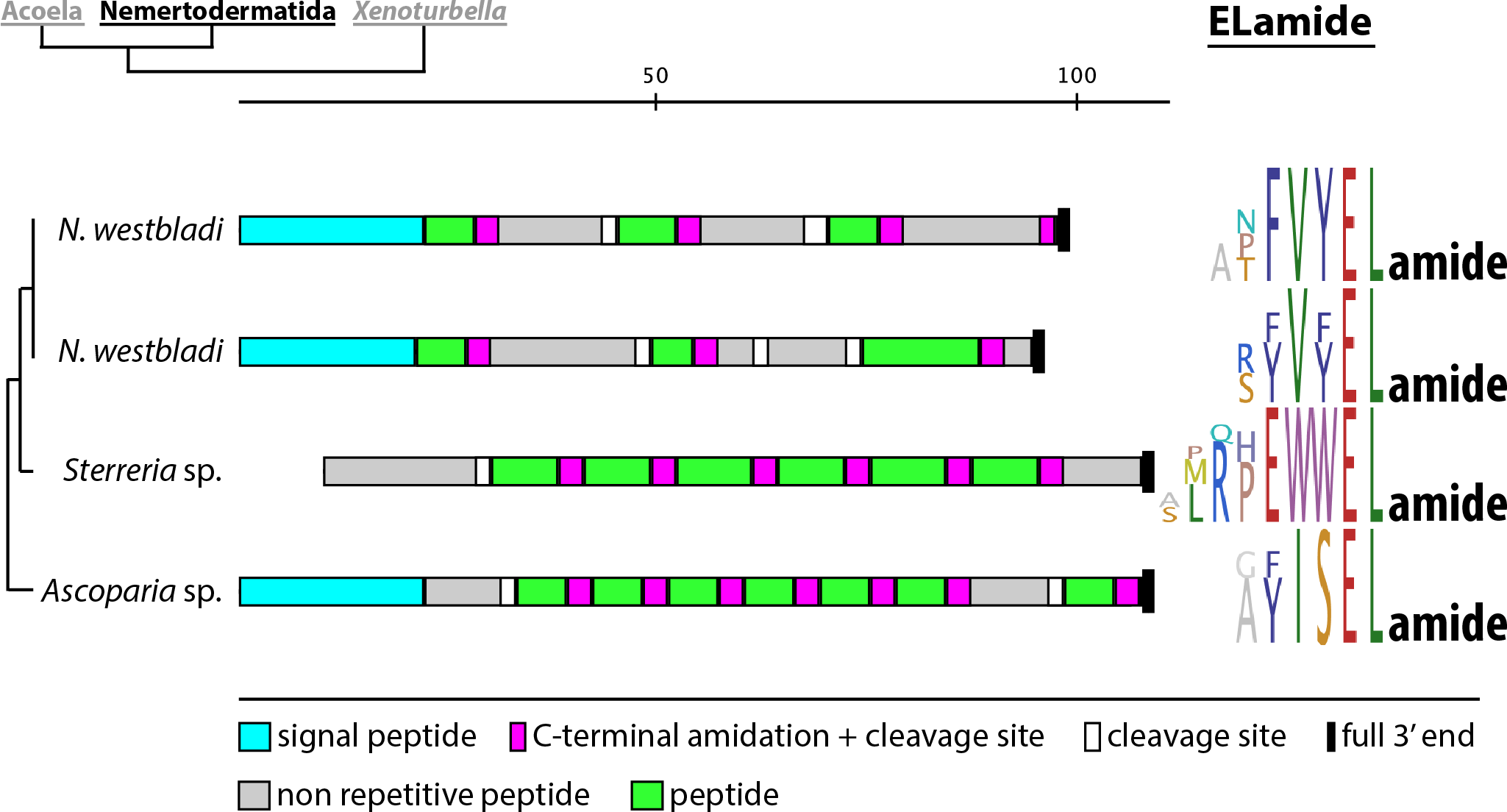
Nemertodermatid ELamide peptides. Precursor scheme and peptide sequence logos of aligned peptides of each precursor. Scale bar above precursor schemes indicates length in amino acids.

#### *LRIGamide peptides* (Figure 25)

Three nemertodermatid transcriptomes contained LRIGamide peptide sequences. *N. westbladi* has 3 paralogs, where two of them only differ slightly in their non-repetitive stretches and the third one has the isoleucine replaced by a valine residue. The *N. westbladi* sequences have 5 copies of the conserved peptide, each separated by a non-conserved part, while the fully recovered sequence of *Ascoparia* sp. has six repeats in two blocks. All paracopies of the peptides are completely identical within the same precursor.

**Figure 25:**
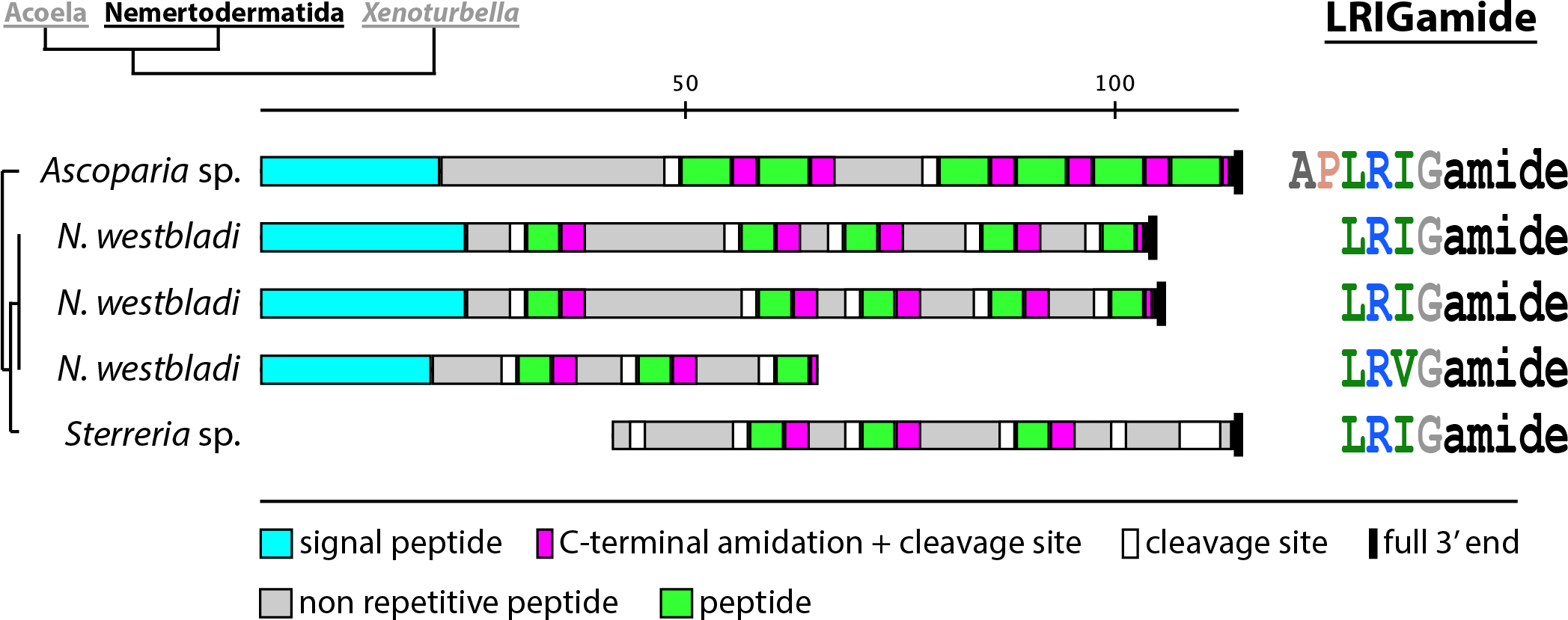
Nemertodermatid LRIGamide peptides. Precursor scheme and highly conserved predicted active peptides. Scale bar above precursor schemes indicates length in amino acids.

### *Xenoturbella* specific MCPs

#### *LRFDIamide peptides* (Figure 26)

The only potential neuropeptides that are specific to the investigated *Xenoturbella* species constitute prepropeptide sequences encoding the pentapeptide LRFDIamide. The precursors of both species are very similar with 4 copies of the peptide in the same arrangement and an overall high conservation of the non-repetitive peptides. The arginine residue of the repetitive peptide is replaced by a lysine in two of the four copies in the *X. bocki* precursor.

**Figure 26:**
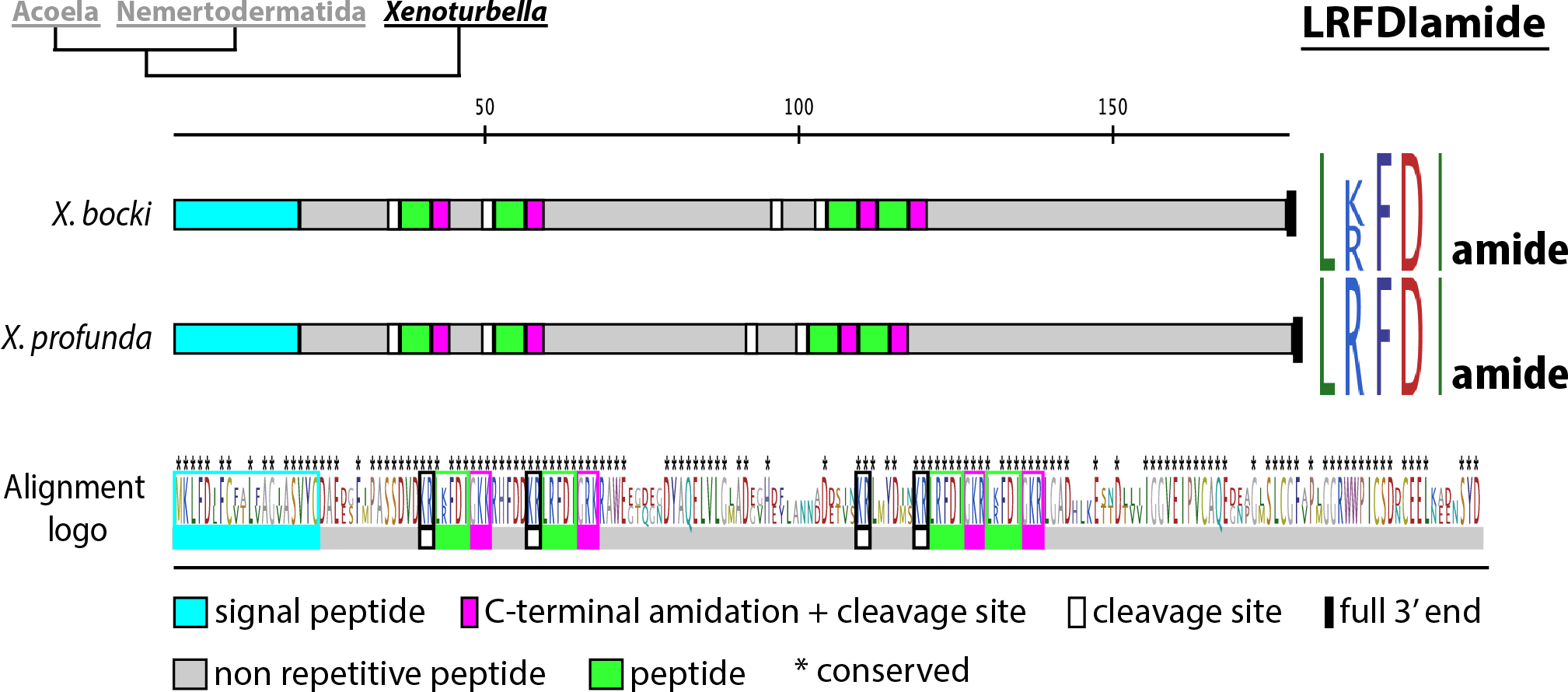
Xenoturbella LRFDIamide peptides. Precursor scheme, peptide sequence logos of aligned peptides of each precursor and p eptide sequence logo of both aligned precursors. Scale bar above precursor schemes indicates length in amino acids. Supplementary Table and Figures

### Single MCPs without orthologs

We found several potential neuropeptide precursors that were only identified in single species, without any indication of orthologues sequences in other transcriptomes. The precursor arrangements and peptide sequence logos of these sequences are shown in Supplementary Figure 6. These precursors exclusively belonged to acoel and nemertodermatid species, whereas no species-specific MCP was identified in either of the two *Xenoturbella* species.

### The xenacoelomorph proneuropeptide complement shows a lack of expected RFamide peptides

We recognized a lack of RFamide peptides in xenacoelomorphs, which was unexpected because neuropeptides with the C-terminal ending RFamide or RYamide are present in nearly all deuterostome and protostome clades as well as in cnidarians [3, 26, 95-98]. The only detected neuropeptides with the C-terminal ending RFamide are a *D. gymnopharyngeus* FxxxFamide, a *C. submaculatum* YLRFamide partial sequence and the *N. westbladi* neuropeptide Y/F-like peptides. The different RFamide-like positive immunostainings with crossreacting antibodies that have been shown in various xenacoelomorph species indicate that there are likely more RFamides, or possibly RYamides present [39, 43, 46-48, 50]. A low representation of certain peptides can be caused by the incompleteness or divergent quality of transcriptomic datasets, as well as a rapid sequence divergence that avoids the detection of orthologs. For a more conclusive picture, genomic data would be preferable to purely transcriptomic data, but the use of several different transcriptomes should -to a certain extend- compensate for incomplete individual transcriptomes and lineage specific divergences and therefore give a good overall picture of the xenacoelomorph neuropeptide complement and clade-specific differences. It has to be kept in mind, that our approach did not let us detect those neuropeptides that only have a single copy on their propeptide and show no clear similarity to known neuropeptides, leading to the assumption that those peptides that crossreacted with the RFamide antibodies are either not multi-copy peptides or their crossreactivity is not based on the exact C-terminal RFamide motif.

### Cnidarian GPCRs

Due to the rather inconclusive suggestions of several cnidarian orthologs with bilaterian neuropeptide GPCRs [25, 27-29], we re-analysed cnidarian neuropeptide receptors that were predicted by automatic annotations from the genomes of *A. digitifera, E. pallida* and *O. faveolata*, as well as sequences that were previously predicted from the *N. vectensis* genome by [25]. The GPCRs of *N. vectensis* that we re-analysed were originally annotated as galanin-like, tachykinin/SIFamide-like, RFamide/neuropeptideFF/GnIH/neuropeptide Y-like and GnRH/vasopressin-like receptors. The automatically annotated sequences of *A. digitifera*, *E. pallida* and *O. faveolata* included neuropeptide FF, neuropeptide Y, orexin, cholecystokinin-, qRFamide-, RYamide, substance-K-, bombesin-, gastrin-, galanin- and somatostatin-GPCRs. The *N. vectensis* receptors show a higher diversity to bilaterian neuropeptide GPCRs, than the automatically annotated GPCR sequences, even though the latter ones were predicted to have a higher diversity according to their annotation (see Supplementary figure 7 for comparision of the predicted and actual relationship of the different cnidarian GPCRs). Our analysis showed, that the *A. digitifera*, *E. pallida* and *O. faveolata* sequences cluster together and do not seem to be direct orthologs of specific types of bilaterian neuropeptide GPCRs (Figure 4 and Supplementary Figure 2). The relationship of the *N. vectensis* GPCRs to certain bilaterian GPCRs is not well supported; however, they show a tendency to be more spread out over the trees. Interestingly, the *N. vectensis* GPCR that was predicted to be related to GnRH/vasopressin also shows (in both phylogenetic analyses) some affinity to the cluster of receptors that contains, beside achatin and neuropeptideS/CCAP receptors, GnRH and vasotocin GPCRs. This stands in contrast to a different analysis, where the authors suggested *N. vectensis* orthologs of several different GPCRs like neuropeptide Y, neuropeptide FF, endothelin or somatostatin, but found no receptor that showed similarity with one of the receptors of the vasotocin/GnRH group [27]. This could indicate that a common ancestor of vasotocin/GnRH/achatin/neuropeptide S GPCRs was already present in the last common ancestor of cnidarians and bilaterians and further diversified into the four bilaterian receptor groups after the cnidarian-bilaterian split.

## Discussion

### Similarities of xenacoelomorph MCPs with clade specific deuterostome and protostome MCPs: ancestral peptides or convergent evolution?

In our analysis, we discovered MCPs that show similar motifs to MCPs that are only known from certain deuterostome or protostome clades and that have been previously considered to be specific to these groups. The similarity to clade-specific neuropeptides might either indicate that these are homologous peptides with ancestral motifs or that these motifs evolved convergently. Several acoel LxFamide that share the SxLHFamide motif and some of the SSxxxFamide peptides show sequence similarity to L-type SALMFamides (Figures 13, 14 and 18) that are so far only known from echinoderms [99-101]. It has been reported that two different antibodies that were raised against echinoderm SALMFamides show immunoreactivity in xenacoelomorphs [49, 51]. While one SALMFamide antibody showed cross-reactivity in different deuterostome and protostome taxa, another one only stained nerves in ambulacrarians, *X. bocki* and three acoel species [49, 51]. These similarities would be in line with the previous notion of an affinity of Xenacoelomorpha to ambulacrarians [30]. Other xenacoelomorph MCPs, however, share certain sequence similarities with neuropeptides of protostomes. One of the protostome clade-specific neuropeptides that show similarity to a certain type of xenacoelomorph multi-copy peptides is the PxFVamide (Figure 16). Peptides with a PxFVamide motif have been described in the molluscs *Mytilus edulis* [102], *Theba pisana* [58], *Lottia gigantea* [92], *Charonia tritonis* [103], Sepia officinalis [104] *Pinctata fucata* and *Crassostrea gigas* [94] and the annelid *Capitella teleta* [57]. All of these peptides are 5-9 amino acids long, are encoded multiple times on their propeptide sequences and are referred to as ‘PxFVamide’ [58, 92, 103] ‘FVamide’ [57] or *‘Mytilus* inhibitory peptide’ [94, 102]. Another kind of xenacoelomorph MCP with a similar motif to a protostome-specific neuropeptide are the ELamides (Figure 24). These peptides, and especially the two *N. westbladi* sequences, show similarity to the arthropod allatostatin A, with their C-terminal sequence [F/Y]x[Y/F]xLamide.

Allatostatin A peptides share the C-terminal ending [Y/F]x[F]GLamide, and is usually encoded several times on its precursor, with slight differences between the single copies that can vary in length between 7-13 amino acids [2, 8, 56, 105-107] [with some exceptions of up to 21 amino acids in a single copy [108]]. Orthologs of the allatostatin A receptors are present in Xenacoelomorpha, including *N. westbladi*, which would make it generally possible to test a receptor activation in order to find out whether the similarities of the peptides are due to convergent or homologous evolution. The receptors of the echinoderm SALMFamides and the trochozoan PxFVamides are to this date unkown, which makes it so far impossible to test homology hypotheses. The general variability of the SALMFamides in echinoderms and the PxFVamides in trochozoans shows that these peptides have a high evolutionary rate within those animal groups. Due to the short length and high diversity of many multi-copy peptides (MCPs), the general possibility of a convergent evolution of similar motifs has to be considered, especially when comparing distant related taxa. RFamide-type neuropeptides for example share a similar C-terminal motif but not all of them are related to each other [1, 2, 26, 96, 98, 109]. A slightly different picture is shown in the achatins that we found in both *Xenoturbella* species and the two nemertodermatids *N. westbladi* and *M. stichopi*. Achatins are especially known from trochozoans, where many of them share the sequence GF[G/A/W]D [55, 57, 58, 94], with a few exceptions where either single [92, 94, 103] or all [104] copies differ in other positions. The xenacoelomorph achatins have identical sequences as the deuterostomes *Saccoglossus* and *Branchiostoma*, and the protostomes *Priapulus* and *Halicryptus*. This could be seen as an indication that the GFGN sequence represents the ancestral achatin sequence or was at least very similar to it.

### The Xenacoelomorph peptidergic systems show an early diversification of bilaterian neuropeptide signaling and clade-specific divergences within Xenacoelomorpha

Our GPCR survey and phylogenetic analysis revealed xenacoelomorph orthologs of at least 20 different types of nephrozoan neuropeptide GPCRs, whereas the previously annotated cnidarian neuropeptide GPCRs did not show specific orthologs of nephrozoan peptidergic systems. This suggests, that many of the extant nephrozoan peptidergic systems have evolved and diverged after the split of bilaterians and cnidarians, early in the ancestral lineage leading to all extant Bilateria. When we compare the bilaterian peptidergic systems and the clade-specific nervous system anatomies of acoels, nemertodermatids and *Xenoturbella*, then we do not find any correlation between the presence of these ancestral bilaterian systems and the presence of a centralized nervous system (Figure 6). The common notion is that the ancestral xenacoelomorph possessed a basiepidermal nerve net, as such a plexus can be found in all three clades, and nerve nets in general are present in non-bilaterian as well as bilaterian animals [36, 37, 40, 42, 45, 110]. Such a basiepidermal nerve net constitutes the entire nervous system of *Xenoturbella*, which has no considerable nerve cord or brain-like condensations at all [43, 44]. In Xenoturbella, we find many of the ancestral bilaterian neuropeptide GPCRs and four of the five ancestral bilaterian ligands, some of which are associated with specific brain regions in different Bilateria, like the hypothalamus of mice or zebra fish and the apical nervous system of *Platynereis dumerilii* [11, 17]. Their presence in *Xenoturbella* shows, that they do not have to be necessarily coupled with the presence of any centralized nervous system with a brain-like structure. Brains and nerve cords can only be found in acoelomorphs, and the ones that are present in extant species do not seem to be homologous to the brains and nerve cords of protostomes or deuterostomes [40, 42, 110]. The brain-like condensations and 1-2 pairs of nerve cords in nemertodermatids are usually situated in a basiepidermal position, which is presumed to be ancestral, whereas the acoel nervous system is often considered to be more derived, and their brains and 2-4 pairs of nerve cords are usually internalized in the body [36-40, 42, 46, 51, 111]. Nemertodermatids as well as acoels possess a variety of clade-specific and species-specific multi-copy peptides, which indicates a stronger subsequent diversification of neuropeptide signaling within Acoelomorpha, Nemertodermatida and Acoela. However, while we find nearly all of the conserved bilaterian neuropeptides and neuropeptide GPCRs in nemertodermatids, acoels show a strong reduction in conserved neuropeptide GPCRs and a complete lack of conserved bilaterian neuropeptide precursors. This is in correlation with the results of a preliminary genomic survey where they generally found a smaller set of conserved GPCRs, including a very low number of secretin type GPCRs, in the acoel *Symsagittifera roscoffensis*, compared to *Xenoturbella* [41]. We also detected no prokineticins and only a single, somewhat derived, insulin-like peptide and glycoprotein hormone related peptide in acoels, all of which indicates that the acoel nervous system is more derived on a neuropeptidergic level as well.

The xenacoelomorph specific neuropeptides and the bilaterian peptidergic systems therefore show to some extend opposite tendencies. The higher variety of clade specific peptides in acoelomorphs correlates with the increased anatomical “nervous system complexity” of nemertodermatids and acoels. Many of the ancestral bilaterian peptidergic systems, however, are present in Nemertodermatida and *Xenoturbella*, but strongly reduced in Acoela. If the ancestral xenacoelomorph nervous system only consisted of a nerve net, and nerve cords and brains evolved only in the acoelomorph lineage, this would indicate that the diversification of the bilaterian peptidergic systems is independent from nerve centralization events. Instead, the evolution of neuropeptide systems might be thus more connected to a general diversification of cell types with novel neural curcuits and the increase in differentiated signaling that is indepenent of synaptic connections. Future studies of the localization of the different neuropeptides and their receptors within the nervous system of different xenacoelomorphs, and especially *Xenoturbella*, could reveal more insights on the evolution of neuropeptides and neurosecretory centers, peptidergic and synaptic connectomes, and their connection to what is often referred to as nervous system complexity.

## Acknowledgements

We want to thank Silke Wahl and Johannes Madlung, for helping with mass spectrometric sample preparations and measurements. We thank the members of S9 lab at Sars, especially Anlaug Furu Boddington for help with collecting and maintaining the fresh material used for the mass spectrometry.

## Supplementary Table and Figures

**Supplementary Table 1:**
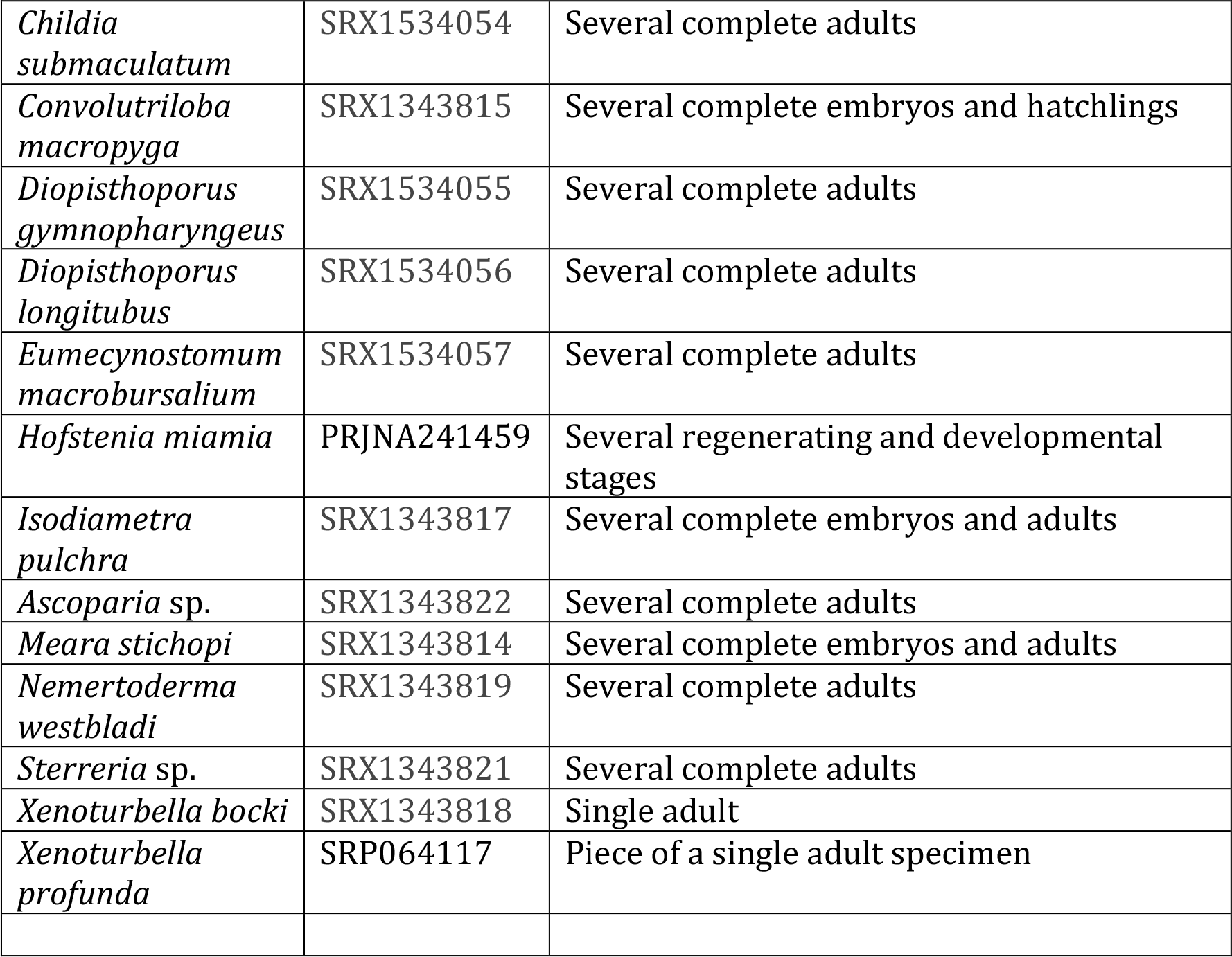
Species, accession number of sequencing data and tissues/stages of RNA source.

**Supplementary Figure 1:**
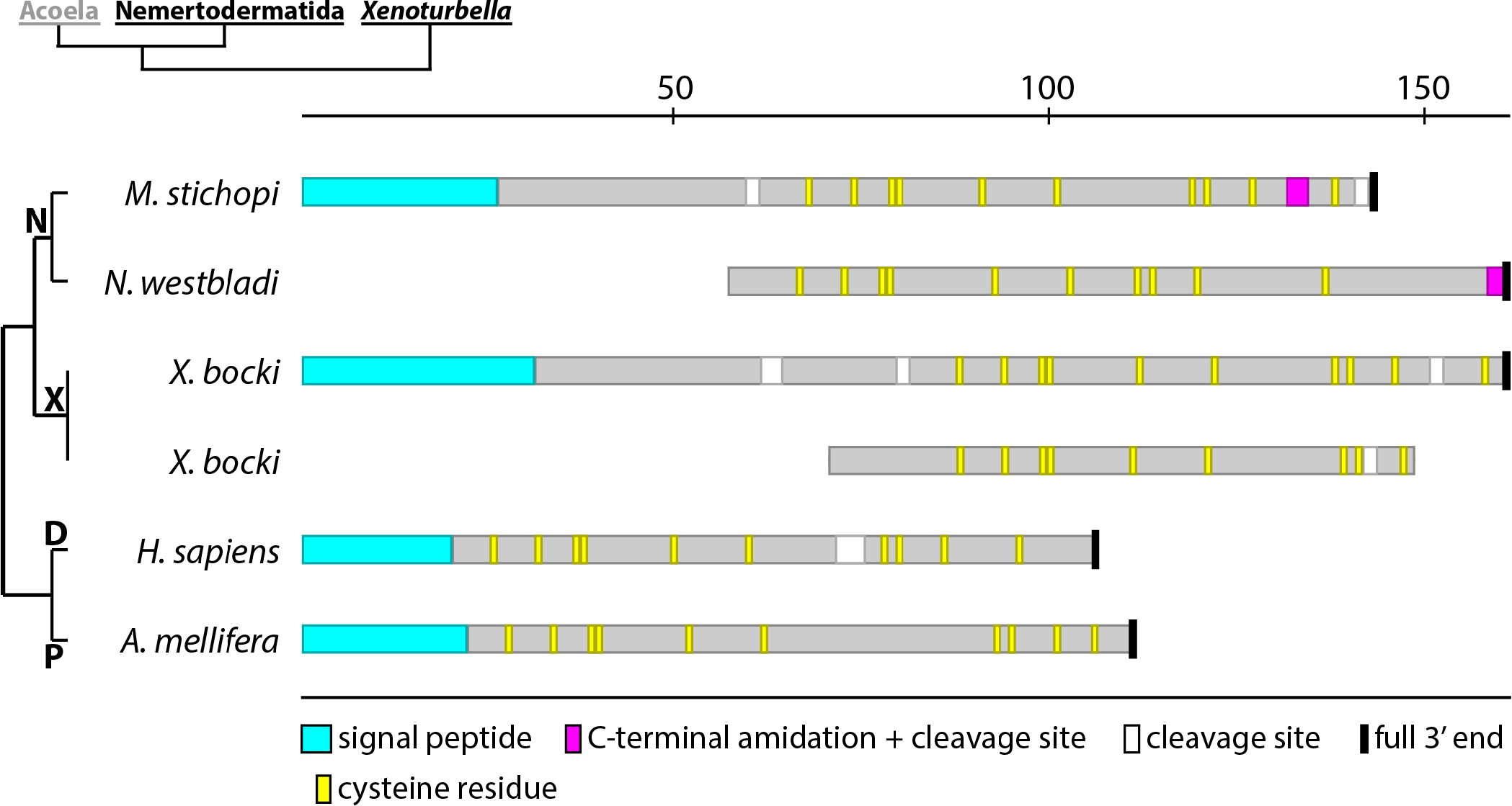
Xenacoelomorph prokineticin like peptide precursor schemes. Homo sapiens prokineticin-1 precursor NP_115790.1, Apis mellifera astakin precursor XP_003250271. Scale bar above precursor schemes indicates length in amino acids. D Deuterostomia, N Nemertodermatida, P Protostomia, *X. Xenoturbella*.

**Supplementary Figure 2:**
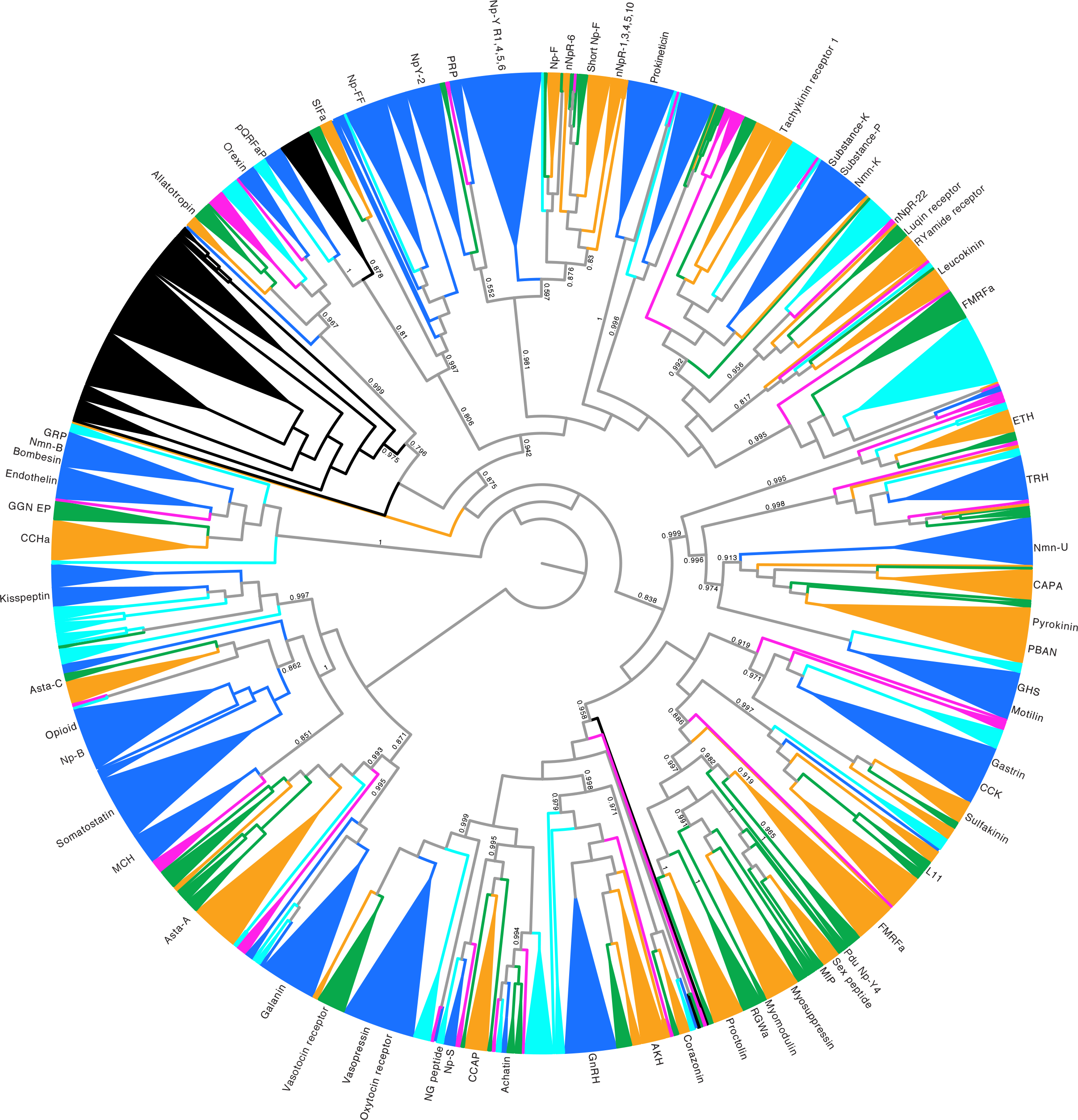
FastTree tree of Rhodopsin type neuropeptide GPCRs. Rhodopsin *beta* GPCRs are rooted against rhodopsin *gamma* GPCRs. Support values (SH-like local support) are shown for the major branches that constitute orthologues receptor types from different animal groups. Branches of homologues receptors that belong to the same animal group with support values of at least 0.8 are combined with the FigTree cartoon-option. *a* amide, *AKH* adipokinetic hormone, *Asta* allatostatin, *CCAP* crustacean cardioaccelatory peptide, *CCK* cholecystokinin, *GGN-EP* GGN excitatory peptide, *ETH* ecdysis triggering hormone, *GHS* growth hormone segretagogue, *GnRH* gonadotropin releasing hormone, *GRP* gastrin releasing peptide, *Np* neuropeptide, *nNpR* nematode neuropeptide receptor, *Nmn* Neuromedin, *L11* elevenin, *MCH* melanin concentrating hormone, *MIP* myoinhibitory peptide, *PBAN* pheromone biosynthesis activating neuropeptide, *pQRFaP* pyroglutaminated RFamide peptide, *PRP* prolactin releasing peptide, *TRH* thryotropin releasing hormone. Colour coding: Magenta = xenacoelomorphs, dark blue = chordates, light blue = ambulacrarians, orange = ecdysozoans, green = spiralians, black = cnidarians.

**Supplementary Figure 3:**
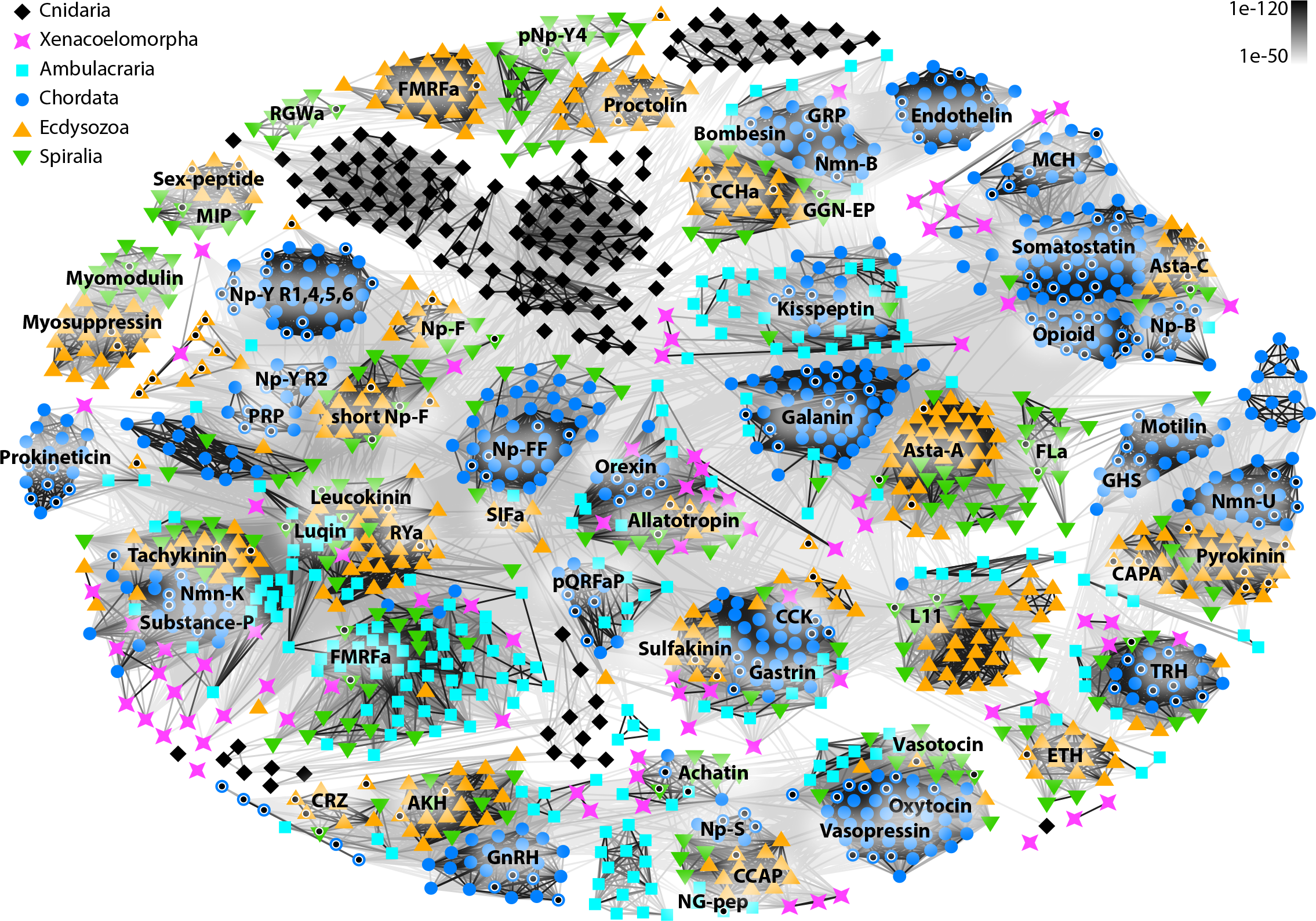
Clustermap of rhodopsin-type neuropeptide GPCRs. Connections are based on blast similarities < 1e-50. A black dot inside a white circle indicates a receptor for which ligand-receptor activation has been shown before. *a* amide, *AKH* adipokinetic hormone, *Asta* allatostatin, *CCAP* crustacean cardioaccelatory peptide, *CCK* cholecystokinin, *GGN-EP* GGN excitatory peptide, *ETH* ecdysis triggering hormone, *GHS* growth hormone segretagogue, *GnRH* gonadotropin releasing hormone, *GRP* gastrin releasing peptide, *Np* neuropeptide, *nNpR* nematode neuropeptide receptor, *Nmn* Neuromedin, *L11* elevenin, *MCH* melanin concentrating hormone, *MIP* myoinhibitory peptide, *PBAN* pheromone biosynthesis activating neuropeptide, *pQRFaP* pyroglutaminated RFamide peptide, *PRP* prolactin releasing peptide, *TRH* thryotropin releasing hormone.

**Supplementary Figure 4:**
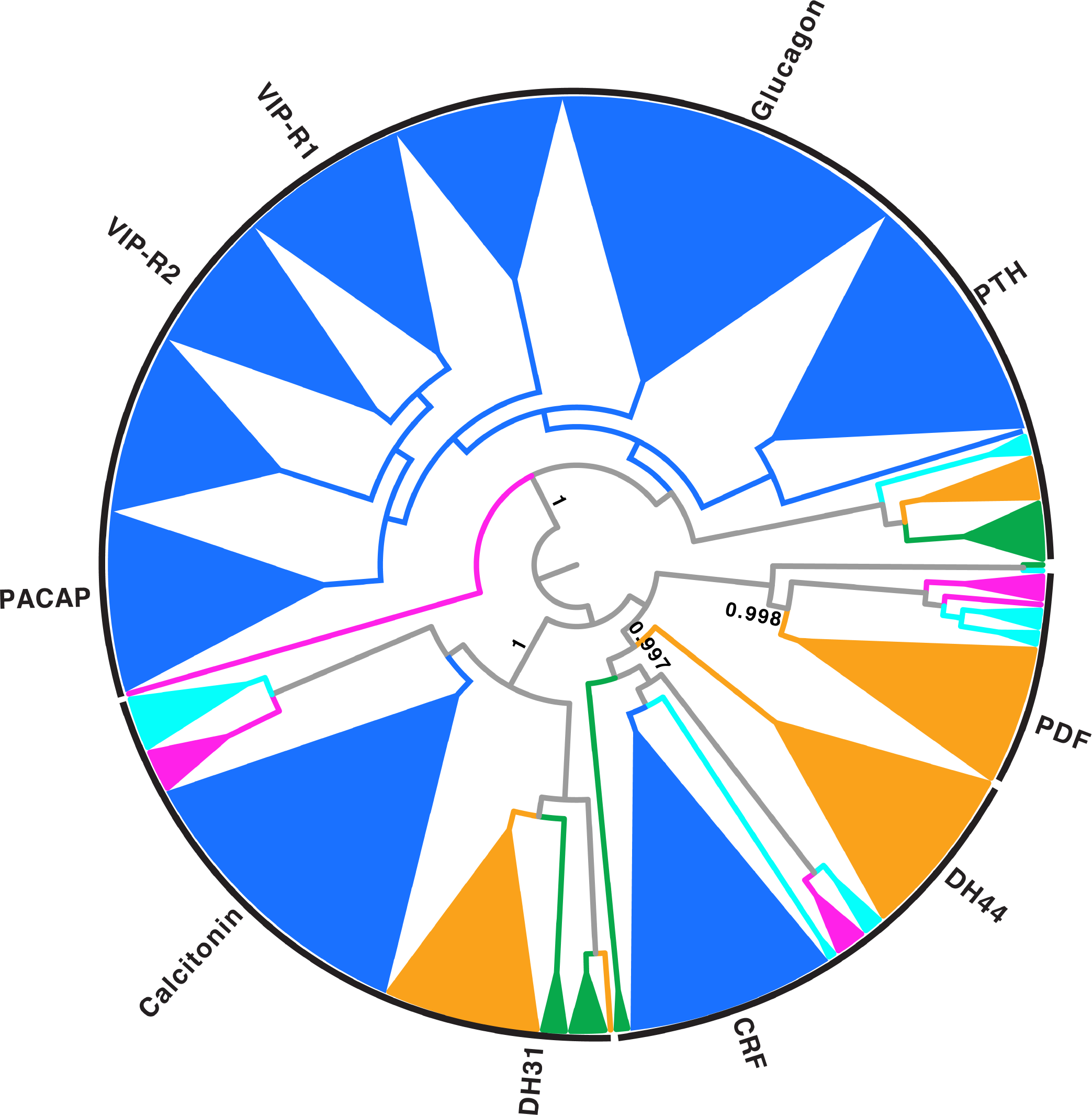
FastTree tree of secretin type neuropeptide GPCRs. Glucagon-related receptor group with 100% support is rooted against the remaining receptors. Support values (SH-like local support) are shown for the major branches that constitute orthologues receptor types from different animal groups. Branches of homologues receptors that belong to the same animal grouping with support values of at least 0.8 are combined with the FigTree cartoon-option. Black lines at the periphery of the tree indicate urbilaterian receptor groups that include xenacoelomorph receptors. *CRF* corticotropin releasing factor, *DH* diuretic hormone, *PDF* pigment dispersing factor, *PTH* parathyroid hormone, *VIP* vasoactive intestinal peptide, *PACAP* pituitary adenylate cyclase-activating polypeptide, -*R* receptor. Colour coding: Magenta = xenacoelomorphs, dark blue = chordates, light blue = ambulacrarians, orange = ecdysozoans, green = spiralians, black = cnidarians.

**Supplementary Figure 5:**
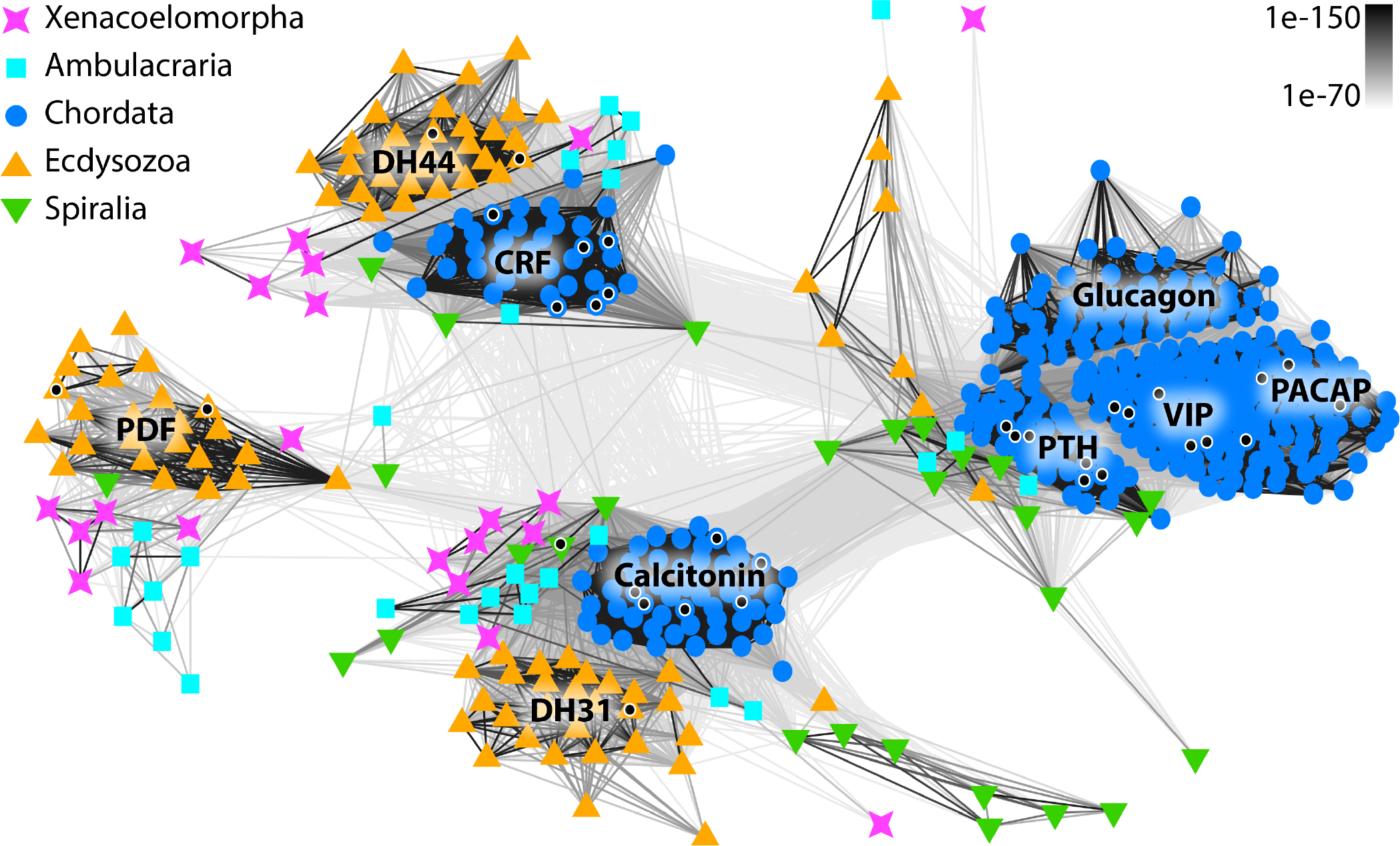
Clustermap of secretin-type neuropeptide GPCRs. Connections are based on blast similarities < 1e-70. A black dot inside a white circle indicates a receptor for which ligand-receptor activation has been shown before. *CRF* corticotropin releasing factor, *DH* diuretic hormone, *PDF* pigment dispersing factor, *PTH* parathyroid hormone, *VIP* vasoactive intestinal peptide, *PACAP* pituitary adenylate cyclase-activating polypeptide.

**Supplementary Figure 6:**
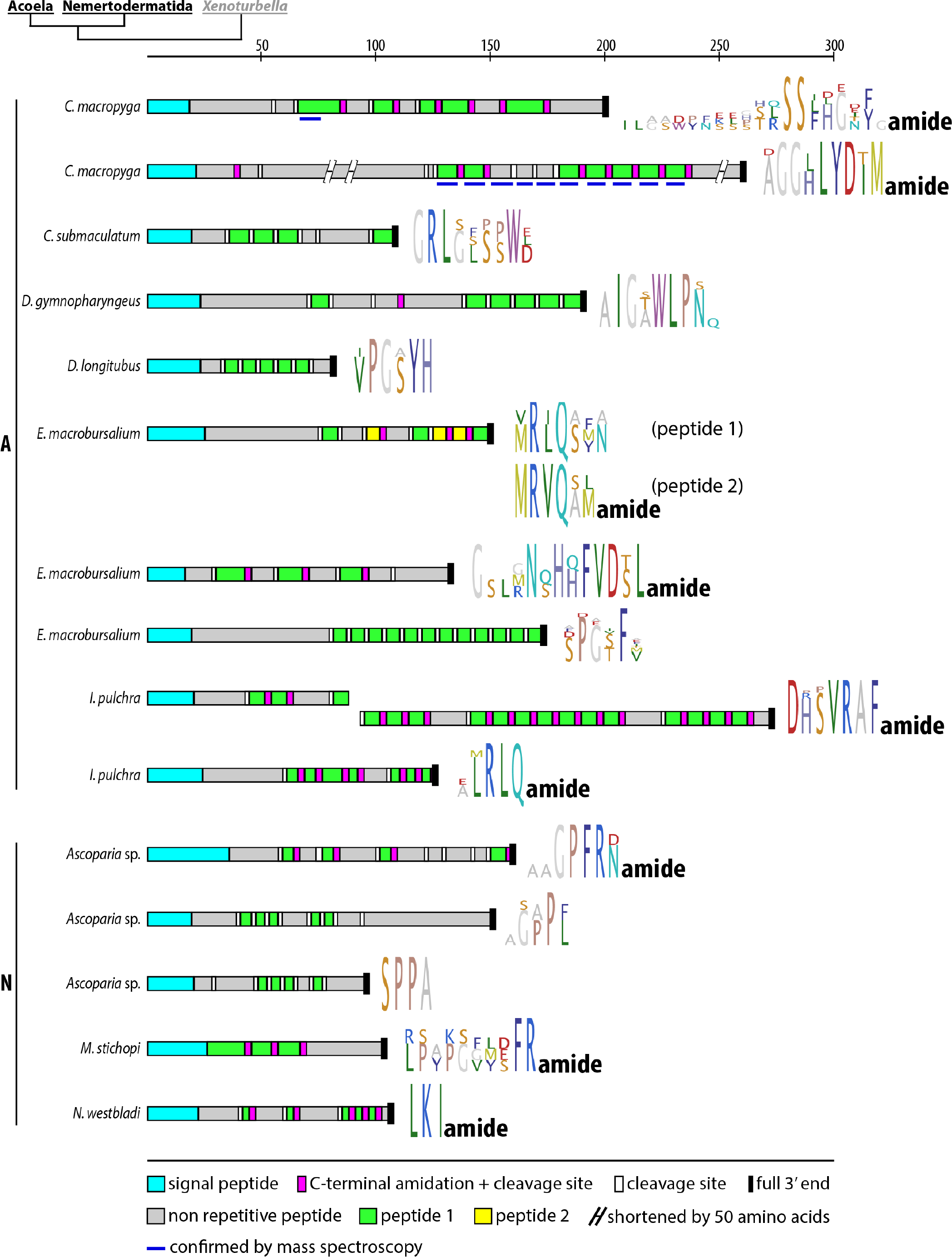
Acoelomorph peptides detected only in single species. Precursor scheme and peptide logos of aligned peptides of each precursor. A Acoela, N Nemertodermatida. Scale bar above precursor schemes indicates length in amino acids.

**Supplementary Figure 7:**
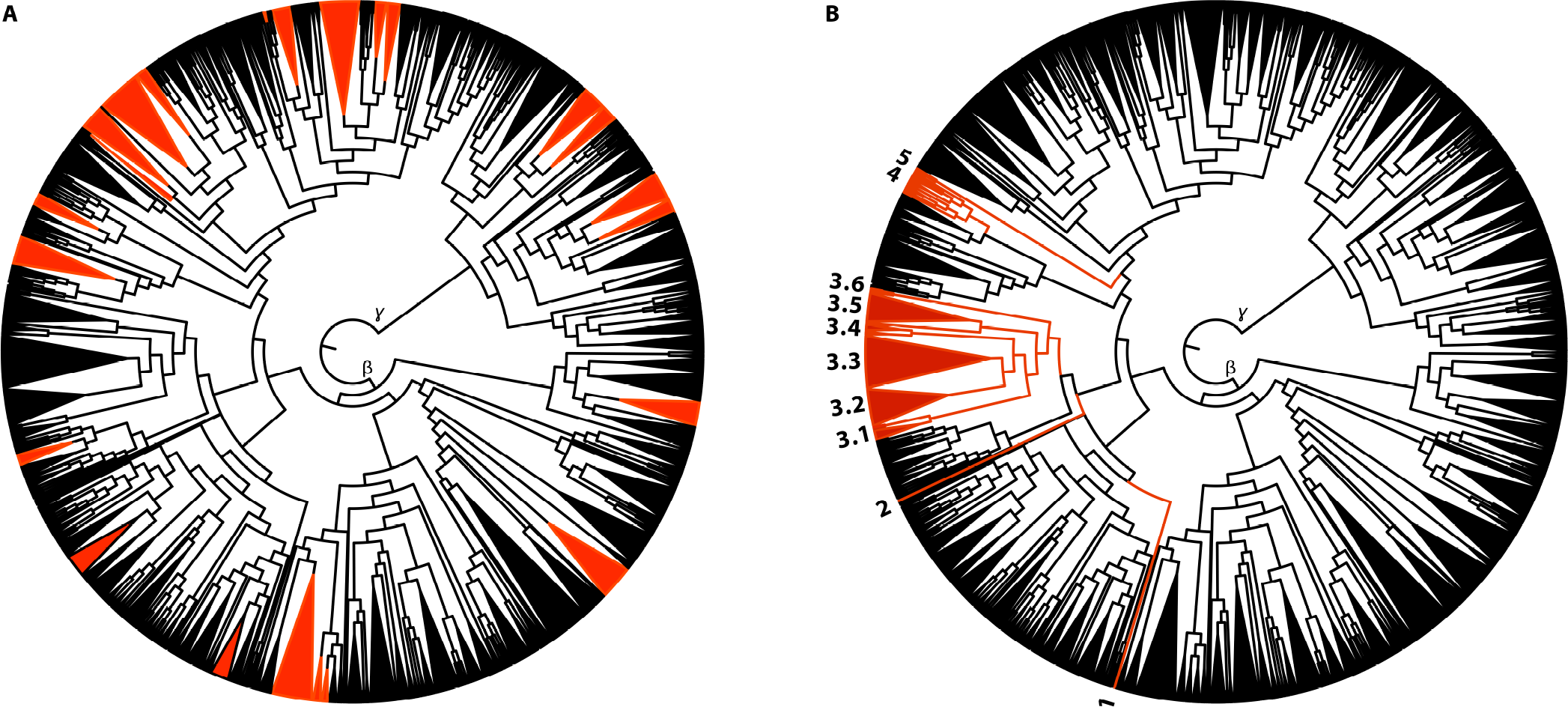
Comparision of sequence distribution of annotated cnidarian GPCRs with the results of the phylogenetic analysis. For annotation of receptor groups see figure 4. A Expected distribution of cnidarian rhodopsin-like GPCRs, based on previous predictions and automated annotations. B Distribution, based on phylogenetic analysis.. Numbered cnidarian groups contain sequences with following annotation: **(1)** *N. vectensis* GnRH/vasotocin; **(2)** *N. vectensis* RFa/Np-FF/GnIH/Np-Y; **(3.1)** *A. digitifera* Np-FF; *E. pallida* orexin, Np-FF; *O. faveolata* galanin, gastrin, Np-FF; **(3.2)** *N. vectensis* galanin; *A. digitifera* bombesin, galanin, orexin, Np-Y R2, Np-Y R6, Np-FF; *E. pallida* bombesin, pQRFaR, Np-FF; *O. faveolata* bombesin, galanin, Np-Y R2, Np-FF; **(3.3)** *N. vectensis* galanin; *A. digitifera* galanin, somatostatin, Np-Y R1, Np-FF; *E. pallida* galanin, somatostatin, substance K, Np-Y R6, Np-FF; *O. faveolata* galanin, substance K, Np-Y R1, Np-Y R6, Np-FF; **(3.4)** *A. digitifera* orexin, cholecystokinin, galanin; *E. pallida* Np-FF, orexin; *O. faveolata* galanin, RYamide; **(3.5)** *N. vectensis* galanin, RFa/Np-FF/GnIH/Np-Y; *A. digitifera* galanin, neuromedin U, orexin, Np-Y 6; *E. pallida* galanin, PRP, Np-Y 2, Np-Y 6; *O. faveolata* galanin, somatostatin, Np-Y 6, Np-Y 2; **(3.6)** *N. vectensis* RFa/Np-FF/GnIH/Np-Y; *E. pallida* cholecystokinin; **(4)** *N. vectensis* tachykinin/SIFamide, RFa/Np-FF/GnIH/Np-Y; **(5)** *N. vectensis* RFa/Np-FF/GnIH/Np-Y.

## References

1 Mirabeau, O., Joly, J. S. 2013 Molecular evolution of peptidergic signaling systems in bilaterians. Proc Natl Acad Sci U S A. 110, E2028-E2037. (10.1073/pnas.1219956110)

2 Jékely, G. 2013 Global view of the evolution and diversity of metazoan neuropeptide signaling. Proc Natl Acad Sci U S A. 110, 8702-8707. (10.1073/pnas.1221833110)

3 Grimmelikhuijzen, C. J. P., Williamson, M., Hansen, G. N. 2002 Neuropeptides in cnidarians. Can J Zool. 80, 1690-1702. (10.1139/Z02-137)

4 Hokfelt, T., Broberger, C., Xu, Z. Q., Sergeyev, V., Ubink, R., Diez, M. 2000 Neuropeptides- an overview. Neuropharmacology. 39, 1337-1356.

5 Roch, G. J., Sherwood, N. M. 2014 Glycoprotein Hormones and Their Receptors Emerged at the Origin of Metazoans. Genome Biol Evol. 6, 1466-1479. (10.1093/gbe/evu118)

6 Stevens, R. C., Cherezov, V., Katritch, V., Abagyan, R., Kuhn, P., Rosen, H., Wuthrich, K. 2013 The GPCR Network: a large-scale collaboration to determine human GPCR structure and function. Nat Rev Drug Discov. 12, 25-34. (10.1038/nrd3859)

7 Frooninckx, L., Van Rompay, L., Temmerman, L., Van Sinay, E., Beets, I., Janssen, T., Husson, S. J., Schoofs, L. 2012 Neuropeptide GPCRs in C. elegans. Front Endocrinol (Lausanne). 3, 167. (10.3389/fendo.2012.00167)

8 Nassel, D. R., Winther, A. M. 2010 Drosophila neuropeptides in regulation of physiology and behavior. Prog Neurobiol. 92, 42-104. (10.1016/j.pneurobio.2010.04.010)

9 Christie, A. E., Skiebe, P., Marder, E. 1995 Matrix of neuromodulators in neurosecretory structures of the crab Cancer borealis. J Exp Biol. 198, 2431-2439.

10 Catak, Z., Aydin, S., Sahin, I., Kuloglu, T., Aksoy, A., Dagli, A. F. 2014 Regulatory neuropeptides (ghrelin, obestatin and nesfatin-1) levels in serum and reproductive tissues of female and male rats with fructose-induced metabolic syndrome. Neuropeptides. 48, 167-177. (10.1016/j.npep.2014.04.002)

11 Williams, E. A., Veraszto, C., Jasek, S., Conzelmann, M., Shahidi, R., Bauknecht, P., Mirabeau, O., Jekely, G. 2017 Synaptic and peptidergic connectome of a neurosecretory centre in the annelid brain. Elife. 6, (10.7554/eLife.26349)

12 Senatore, A., Reese, T. S., Smith, C. L. 2017 Neuropeptidergic integration of behavior in Trichoplax adhaerens, an animal without synapses. J Exp Biol. 220, 3381-3390. (10.1242/jeb.162396)

13 Diao, F. C., Elliott, A. D., Diao, F. Q., Shah, S., White, B. H. 2017 Neuromodulatory connectivity defines the structure of a behavioral neural network. Elife. 6, (10.7554/eLife.29797)

14 Jekely, G., Melzer, S., Beets, I., Kadow, I. C. G., Koene, J., Haddad, S., Holden-Dye, L. 2018 The long and the short of it - a perspective on peptidergic regulation of circuits and behaviour. J Exp Biol. 221, (10.1242/jeb.166710)

15 Conzelmann, M., Jékely, G. 2012 Antibodies against conserved amidated neuropeptide epitopes enrich the comparative neurobiology toolbox. Evodevo. 3, (10.1186/2041-9139-3-23)

16 Kerbl, A., Conzelmann, M., Jekely, G., Worsaae, K. 2017 High diversity in neuropeptide immunoreactivity patterns among three closely related species of Dinophilidae (Annelida). J Comp Neurol. 525, 3596-3635. (10.1002/cne.24289)

17 Tessmar-Raible, K., Raible, F., Christodoulou, F., Guy, K., Rembold, M., Hausen, H., Arendt, D. 2007 Conserved sensory-neurosecretory cell types in annelid and fish forebrain: insights into hypothalamus evolution. Cell. 129, 1389-1400. (10.1016/j.cell.2007.04.041)

18 Grimmelikhuijzen, C. J., Hauser, F. 2012 Mini-review: the evolution of neuropeptide signaling. Regul Pept. 177 Suppl, S6–9. (10.1016/j.regpep.2012.05.001)

19 Semmens, D. C., Beets, I., Rowe, M. L., Blowes, L. M., Oliveri, P., Elphick, M. R. 2015 Discovery of sea urchin NGFFFamide receptor unites a bilaterian neuropeptide family. Open Biol. 5, 150030. (10.1098/rsob.150030)

20 Van Sinay, E., Mirabeau, O., Depuydt, G., Van Hiel, M. B., Peymen, K., Watteyne, J., Zels, S., Schoofs, L., Beets, I. 2017 Evolutionarily conserved TRH neuropeptide pathway regulates growth in Caenorhabditis elegans. Proc Natl Acad Sci U S A. 114, E4065-E4074. (10.1073/pnas.1617392114)

21 Janssen, T., Meelkop, E., Lindemans, M., Verstraelen, K., Husson, S. J., Temmerman, L., Nachman, R. J., Schoofs, L. 2008 Discovery of a cholecystokinin-gastrin-like signaling system in nematodes. Endocrinology. 149, 2826-2839. (10.1210/en.2007-1772)

22 Zandawala, M., Tian, S., Elphick, M. R. 2017 The evolution and nomenclature of GnRH-type and corazonin-type neuropeptide signaling systems. Gen Comp Endocrinol. (10.1016/j.ygcen.2017.06.007)

23 Tian, S., Zandawala, M., Beets, I., Baytemur, E., Slade, S. E., Scrivens, J. H., Elphick, M. R. 2016 Urbilaterian origin of paralogous GnRH and corazonin neuropeptide signalling pathways. Sci Rep. 6, 28788. (10.1038/srep28788)

24 Elphick, M. R., Mirabeau, O., Larhammar, D. 2018 Evolution of neuropeptide signalling systems. J Exp Biol. 221, (10.1242/jeb.151092)

25 Anctil, M. 2009 Chemical transmission in the sea anemone Nematostella vectensis: A genomic perspective. Comp Biochem Physiol Part D Genomics Proteomics. 4, 268-289. (10.1016/j.cbd.2009.07.001)

26 Walker, R. J., Papaioannou, S., Holden-Dye, L. 2009 A review of FMRFamide- and RFamide-like peptides in metazoa. Invert Neurosci. 9, 111-153. (10.1007/s10158-010-0097-7)

27 Krishnan, A., Schioth, H. B. 2015 The role of G protein-coupled receptors in the early evolution of neurotransmission and the nervous system. J Exp Biol. 218, 562-571. (10.1242/jeb.110312)

28 Alzugaray, M. E., Hernandez-Martinez, S., Ronderos, J. R. 2016 Somatostatin signaling system as an ancestral mechanism: Myoregulatory activity of an Allatostatin-C peptide in Hydra. Peptides. 82, 67-75. (10.1016/j.peptides.2016.05.011)

29 Alzugaray, M. E., Adami, M. L., Diambra, L. A., Hernandez-Martinez, S., Damborenea, C., Noriega, F. G., Ronderos, J. R. 2013 Allatotropin: An Ancestral Myotropic Neuropeptide Involved in Feeding. Plos One. 8, (10.1371/journal.pone.0077520)

30 Philippe, H., Brinkmann, H., Copley, R. R., Moroz, L. L., Nakano, H., Poustka, A. J., Wallberg, A., Peterson, K. J., Telford, M. J. 2011 Acoelomorph flatworms are deuterostomes related to Xenoturbella. Nature. 470, 255-258. (10.1038/nature09676)

31 Cannon, J. T., Vellutini, B. C., Smith, J., Ronquist, F. R., Jondelius, U., Hejnol, A. 2016 Xenacoelomorpha is the sister group to Nephrozoa. Nature. 530, 89-+. (10.1038/nature16520)

32 Rouse, G. W., Wilson, N. G., Carvajal, J. I., Vrijenhoek, R. C. 2016 New deep-sea species of Xenoturbella and the position of Xenacoelomorpha. Nature. 530, 94-97. (10.1038/nature16545)

33 Srivastava, M., Mazza-Curll, K. L., van Wolfswinkel, J. C., Reddien, P. W. 2014 Whole-body acoel regeneration is controlled by Wnt and Bmp-Admp signaling. Curr Biol. 24, 1107-1113. (10.1016/j.cub.2014.03.042)

34 Hejnol, A., Obst, M., Stamatakis, A., Ott, M., Rouse, G. W., Edgecombe, G. D., Martinez, P., Baguna, J., Bailly, X., Jondelius, U., et al. 2009 Assessing the root of bilaterian animals with scalable phylogenomic methods. Proc Biol Sci. 276, 4261-4270. (10.1098/rspb.2009.0896)

35 Ruiz-Trillo, I., Riutort, M., Littlewood, D. T., Herniou, E. A., Baguna, J. 1999 Acoel flatworms: earliest extant bilaterian Metazoans, not members of Platyhelminthes. Science. 283, 1919-1923.

36 Raikova, O. I., Meyer-Wachsmuth, I., Jondelius, U. 2016 The plastic nervous system of Nemertodermatida. Org Divers Evol. 16, 85-104. (10.1007/s13127-015-0248-0)

37 Hejnol, A. 2015 Acoelomorpha. In Structure and Evolution of Invertebrate Nervous Systems. (eds. A. Schmidt-Rhaesa, S. Harzsch, G. Purschke), pp. 56-61: Oxford University Press.

38 Hejnol, A., Pang, K. 2016 Xenacoelomorpha’s significance for understanding bilaterian evolution. Curr Opin Genet Dev. 39, 48-54. (10.1016/j.gde.2016.05.019)

39 Achatz, J. G., Martinez, P. 2012 The nervous system of Isodiametra pulchra (Acoela) with a discussion on the neuroanatomy of the Xenacoelomorpha and its evolutionary implications. Front Zool. 9, 27. (10.1186/1742-9994-9-27)

40 Haszprunar, G. 2016 Review of data for a morphological look on Xenacoelomorpha (Bilateria incertae sedis). Org Divers Evol. 16, 363-389. (10.1007/s13127-015-0249-z)

41 Perea-Atienza, E., Gavilan, B., Chiodin, M., Abril, J. F., Hoff, K. J., Poustka, A. J., Martinez, P. 2015 The nervous system of Xenacoelomorpha: a genomic perspective. J Exp Biol. 218, 618-628. (10.1242/jeb.110379)

42 Gavilan, B., Perea-Atienza, E., Martinez, P. 2016 Xenacoelomorpha: a case of independent nervous system centralization? Philos Trans R Soc Lond B Biol Sci. 371, 20150039. (10.1098/rstb.2015.0039)

43 Raikova, O. I., Reuter, M., Jondelius, U., Gustafsson, M. K. S. 2000 An immunocytochemical and ultrastructural study of the nervous and muscular systems of Xenoturbella westbladi (Bilateria inc. sed.). Zoomorphology. 120, 107-118. (DOI 10.1007/s004350000028)

44 Stach, T. 2015 Xenoturbella. In Structure and Evolution of Invertebrate Nervous Systems. (eds. A. Schmidt-Rhaesa, S. Harzsch, G. Purschke), pp. 62-66: Oxford University Press.

45 Hejnol, A., Rentzsch, F. 2015 Neural nets. Curr Biol. 25, R782-786. (10.1016/j.cub.2015.08.001)

46 Raikova, O. I., Reuter, M., Gustafsson, M. K., Maule, A. G., Halton, D. W., Jondelius, U. 2004 Basiepidermal nervous system in Nemertoderma westbladi (Nemertodermatida): GYIRFamide immunoreactivity. Zoology (Jena). 107, 75-86. (10.1016/j.zool.2003.12.002)

47 Børve, A., Hejnol, A. 2014 Development and juvenile anatomy of the nemertodermatid Meara stichopi (Bock) Westblad 1949 (Acoelomorpha). Front Zool. 11, 50. (10.1186/1742-9994-11-50)

48 Semmler, H., Chiodin, M., Bailly, X., Martinez, P., Wanninger, A. 2010 Steps towards a centralized nervous system in basal bilaterians: insights from neurogenesis of the acoel Symsagittifera roscoffensis. Dev Growth Differ. 52, 701-713. (10.1111/j.1440-169X.2010.01207.x)

49 Stach, T., Dupont, S., Israelson, O., Fauville, G., Nakano, H., Kanneby, T., Thorndyke, M. 2005 Nerve cells of Xenoturbella bocki (phylum uncertain) and Harrimania kupfferi (Enteropneusta) are positively immunoreactive to antibodies raised against echinoderm neuropeptides. J Mar Biol Assoc Uk. 85, 1519-1524. (Doi 10.1017/S0025315405012725)

50 Reuter, M., Raikova, O. I., Jondelius, U., Gustafsson, M. K. S., Maule, A. G., Halton, D. W. 2001 Organisation of the nervous system in the Acoela: an immunocytochemical study. Tissue Cell. 33, 119-128.

51 Dittmann, I. L., Zauchner, T., Nevard, L. M., Telford, M. J., Egger, B. 2018 SALMFamide2 and serotonin immunoreactivity in the nervous system of some acoels (Xenacoelomorpha). Journal of Morphology. 1-9. (10.1002/jmor.20794)

52 Kotikova, E. A., Raiikova, O. I. 2008 Architectonics of the central nervous system in Acoela, Plathelminthes, and Rotifera. Zh Evol Biokhim Fiziol. 44, 83-93.

53 Thiel, D., Bauknecht, P., Jekely, G., Hejnol, A. 2017 An ancient FMRFamide-related peptide-receptor pair induces defence behaviour in a brachiopod larva. Open Biol. 7, (10.1098/rsob.170136)

54 Semmens, D. C., Mirabeau, O., Moghul, I., Pancholi, M. R., Wurm, Y., Elphick, M. R. 2016 Transcriptomic identification of starfish neuropeptide precursors yields new insights into neuropeptide evolution. Open Biol. 6, (10.1098/rsob.150224)

55 Conzelmann, M., Williams, E. A., Krug, K., Franz-Wachtel, M., Macek, B., Jekely, G. 2013 The neuropeptide complement of the marine annelid Platynereis dumerilii. Bmc Genomics. 14, 906. (10.1186/1471-2164-14-906)

56 Hauser, F., Neupert, S., Williamson, M., Predel, R., Tanaka, Y., Grimmelikhuijzen, C. J. 2010 Genomics and peptidomics of neuropeptides and protein hormones present in the parasitic wasp Nasonia vitripennis. J Proteome Res. 9, 5296-5310. (10.1021/pr100570j)

57 Veenstra, J. A. 2011 Neuropeptide evolution: neurohormones and neuropeptides predicted from the genomes of Capitella teleta and Helobdella robusta. Gen Comp Endocrinol. 171, 160-175. (10.1016/j.ygcen.2011.01.005)

58 Adamson, K. J., Wang, T., Zhao, M., Bell, F., Kuballa, A. V., Storey, K. B., Cummins, S. F. 2015 Molecular insights into land snail neuropeptides through transcriptome and comparative gene analysis. Bmc Genomics. 16, 308. (10.1186/s12864-015-1510-8)

59 Bauknecht, P., Jékely, G. 2015 Large-Scale Combinatorial Deorphanization of Platynereis Neuropeptide GPCRs. Cell Rep. 12, 684-693. (10.1016/j.celrep.2015.06.052)

60 Suwansa-Ard, S., Chaiyamoon, A., Talarovicova, A., Tinikul, R., Tinikul, Y., Poomtong, T., Elphick, M. R., Cummins, S. F., Sobhon, P. 2017 Transcriptomic discovery and comparative analysis of neuropeptide precursors in sea cucumbers (Holothuroidea). Peptides. (10.1016/j.peptides.2017.10.008)

61 Frickey, T., Lupas, A. 2004 CLANS: a Java application for visualizing protein families based on pairwise similarity. Bioinformatics. 20, 3702-3704. (10.1093/bioinformatics/bth444)

62 Martin-Duran, J. M., Ryan, J. F., Vellutini, B., Pang, K., Hejnol, A. 2017 Increased taxon sampling reveals thousands of hidden orthologs in flatworms. Genome Res. (10.1101/gr.216226.116)

63 Larkin, M. A., Blackshields, G., Brown, N. P., Chenna, R., McGettigan, P. A., McWilliam, H., Valentin, F., Wallace, I. M., Wilm, A., Lopez, R., et al. 2007 Clustal W and Clustal X version 2.0. Bioinformatics. 23, 2947-2948. (10.1093/bioinformatics/btm404)

64 Capella-Gutierrez, S., Silla-Martinez, J. M., Gabaldon, T. 2009 trimAl: a tool for automated alignment trimming in large-scale phylogenetic analyses. Bioinformatics. 25, 1972-1973. (10.1093/bioinformatics/btp348)

65 Price, M. N., Dehal, P. S., Arkin, A. P. 2010 FastTree 2--approximately maximum-likelihood trees for large alignments. PLoS One. 5, e9490. (10.1371/journal.pone.0009490)

66 Zhang, Y. Z., Shen, H. B. 2017 Signal-3L 2.0: A Hierarchical Mixture Model for Enhancing Protein Signal Peptide Prediction by Incorporating Residue-Domain Cross-Level Features. J Chem Inf Model. 57, 988-999. (10.1021/acs.jcim.6b00484)

67 Petersen, T. N., Brunak, S., von Heijne, G., Nielsen, H. 2011 SignalP 4.0: discriminating signal peptides from transmembrane regions. Nat Methods. 8, 785-786. (10.1038/nmeth.1701)

68 Southey, B. R., Amare, A., Zimmerman, T. A., Rodriguez-Zas, S. L., Sweedler, J. V. 2006 NeuroPred: a tool to predict cleavage sites in neuropeptide precursors and provide the masses of the resulting peptides. Nucleic Acids Res. 34, W267-272. (10.1093/nar/gkl161)

69 Jondelius, U., Wallberg, A., Hooge, M., Raikova, O. I. 2011 How the Worm Got its Pharynx: Phylogeny, Classification and Bayesian Assessment of Character Evolution in Acoela. Systematic Biology. 60, 845-871. (10.1093/sysbio/syr073)

70 Borchert, N., Dieterich, C., Krug, K., Schutz, W., Jung, S., Nordheim, A., Sommer, R. J., Macek, B. 2010 Proteogenomics of Pristionchus pacificus reveals distinct proteome structure of nematode models. Genome Res. 20, 837-846. (10.1101/gr.103119.109)

71 Rappsilber, J., Mann, M., Ishihama, Y. 2007 Protocol for micro-purification, enrichment, prefractionation and storage of peptides for proteomics using StageTips. Nat Protoc. 2, 1896-1906. (10.1038/nprot.2007.261)

72 Cox, J., Mann, M. 2008 MaxQuant enables high peptide identification rates, individualized p.p.b.- range mass accuracies and proteome-wide protein quantification. Nat Biotechnol. 26, 1367-1372. (10.1038/nbt.1511)

73 Cox, J., Neuhauser, N., Michalski, A., Scheltema, R. A., Olsen, J. V., Mann, M. 2011 Andromeda: A Peptide Search Engine Integrated into the MaxQuant Environment. J Proteome Res. 10, 1794-1805. (10.1021/pr101065j)

74 Elias, J. E., Gygi, S. P. 2007 Target-decoy search strategy for increased confidence in large-scale protein identifications by mass spectrometry. Nature Methods. 4, 207-214. (10.1038/nmeth1019)

75 Luo, C. W., Dewey, E. M., Sudo, S., Ewer, J., Hsu, S. Y., Honegger, H. W., Hsueh, A. J. 2005 Bursicon, the insect cuticle-hardening hormone, is a heterodimeric cystine knot protein that activates G protein-coupled receptor LGR2. Proc Natl Acad Sci U S A. 102, 2820-2825. (10.1073/pnas.0409916102)

76 Nikitin, M. 2015 Bioinformatic prediction of Trichoplax adhaerens regulatory peptides. Gen Comp Endocr. 212, 145-155. (10.1016/j.ygcen.2014.03.049)

77 Steinmetz, P., Aman, A., Kraus, J., Technau, U. 2017 Gut-like ectodermal tissue in a sea anemone challenges germ layer homology. Mech Develop. 145, S111-S111. (10.1016/j.mod.2017.04.295)

78 Chan, S. J., Steiner, D. F. 2000 Insulin through the ages: Phylogeny of a growth promoting and metabolic regulatory hormone. Am Zool. 40, 213-222.

79 Claeys, I., Simonet, G., Poels, J., Van Loy, T., Vercammen, L., De Loof, A., Vanden Broeck, J. 2002 Insulin-related peptides and their conserved signal transduction pathway. Peptides. 23, 807-816.

80 Mizoguchi, A., Okamoto, N. 2013 Insulin-like and IGF-like peptides in the silkmoth Bombyx mori: discovery, structure, secretion, and function. Front Physiol. 4, 217. (10.3389/fphys.2013.00217)

81 Steele, R. E., Lieu, P., Mai, N. H., Shenk, M. A., Sarras, M. P. 1996 Response to insulin and the expression pattern of a gene encoding an insulin receptor homologue suggest a role for an insulin-like molecule in regulating growth and patterning in Hydra. Dev Genes Evol. 206, 247-259. (DOI 10.1007/s004270050050)

82 Liutkeviciute, Z., Koehbach, J., Eder, T., Gil-Mansilla, E., Gruber, C. W. 2016 Global map of oxytocin/vasopressin-like neuropeptide signalling in insects. Sci Rep-Uk. 6, (10.1038/srep39177)

83 Hauser, F., Grimmelikhuijzen, C. J. P. 2014 Evolution of the AKH/corazonin/ACP/GnRH receptor superfamily and their ligands in the Protostomia. Gen Comp Endocrinol. 209, 35-49. (10.1016/j.ygcen.2014.07.009)

84 Hansen, K. K., Stafflinger, E., Schneider, M., Hauser, F., Cazzamali, G., Williamson, M., Kollmann, M., Schachtner, J., Grimmelikhuijzen, C. J. 2010 Discovery of a novel insect neuropeptide signaling system closely related to the insect adipokinetic hormone and corazonin hormonal systems. J Biol Chem. 285, 10736-10747. (10.1074/jbc.M109.045369)

85 Lindemans, M., Janssen, T., Beets, I., Temmerman, L., Meelkop, E., Schoofs, L. 2011 Gonadotropin-releasing hormone and adipokinetic hormone signaling systems share a common evolutionary origin. Front Endocrinol (Lausanne). 2, 16. (10.3389/fendo.2011.00016)

86 Roch, G. J., Busby, E. R., Sherwood, N. M. 2011 Evolution of GnRH: diving deeper. Gen Comp Endocrinol. 171, 1-16. (10.1016/j.ygcen.2010.12.014)

87 Lindemans, M., Liu, F., Janssen, T., Husson, S. J., Mertens, I., Gade, G., Schoofs, L. 2009 Adipokinetic hormone signaling through the gonadotropin-releasing hormone receptor modulates egg-laying in Caenorhabditis elegans. Proc Natl Acad Sci U S A. 106, 1642-1647. (10.1073/pnas.0809881106)

88 Roch, G. J., Tello, J. A., Sherwood, N. M. 2014 At the transition from invertebrates to vertebrates, a novel GnRH-like peptide emerges in amphioxus. Mol Biol Evol. 31, 765-778. (10.1093/molbev/mst269)

89 Li, S., Hauser, F., Skadborg, S. K., Nielsen, S. V., Kirketerp-Moller, N., Grimmelikhuijzen, C. J. 2016 Adipokinetic hormones and their G protein-coupled receptors emerged in Lophotrochozoa. Sci Rep. 6, 32789. (10.1038/srep32789)

90 Kamatani, Y., Minakata, H., Kenny, P. T., Iwashita, T., Watanabe, K., Funase, K., Sun, X. P., Yongsiri, A., Kim, K. H., Novales-Li, P., et al. 1989 Achatin-I, an endogenous neuroexcitatory tetrapeptide from Achatina fulica Ferussac containing a D-amino acid residue. Biochem Biophys Res Commun. 160, 1015-1020.

91 Nassel, D. R., Wegener, C. 2011 A comparative review of short and long neuropeptide F signaling in invertebrates: Any similarities to vertebrate neuropeptide Y signaling? Peptides. 32, 1335-1355. (10.1016/j.peptides.2011.03.013)

92 Veenstra, J. A. 2010 Neurohormones and neuropeptides encoded by the genome of Lottia gigantea, with reference to other mollusks and insects. Gen Comp Endocrinol. 167, 86-103. (10.1016/j.ygcen.2010.02.010)

93 McVeigh, P., Mair, G. R., Atkinson, L., Ladurner, P., Zamanian, M., Novozhilova, E., Marks, N. J., Day, T. A., Maule, A. G. 2009 Discovery of multiple neuropeptide families in the phylum Platyhelminthes. Int J Parasitol. 39, 1243-1252. (10.1016/j.ijpara.2009.03.005)

94 Stewart, M. J., Favrel, P., Rotgans, B. A., Wang, T. F., Zhao, M., Sohail, M., O’Connor, W. A., Elizur, A., Henry, J., Cummins, S. F. 2014 Neuropeptides encoded by the genomes of the Akoya pearl oyster Pinctata fucata and Pacific oyster Crassostrea gigas: a bioinformatic and peptidomic survey. Bmc Genomics. 15, (10.1186/1471-2164-15-840)

95 Dockray, G. J. 2004 The expanding family of -RFamide peptidesand their effects on feeding behaviour. Exp Physiol. 89, 229-235. (10.1113/expphysiol.2004.027169)

96 Espinoza, E., Carrigan, M., Thomas, S. G., Shaw, G., Edison, A. S. 2000 A Statistical View of FMRFamide Neuropeptide Diversity. Mol Neurobiol. 21, 35-56. (10.1385/MN:21:1-2:035)

97 Orchard, I., Lange, A. B., Bendena, W. G. 2001 FMRFamide-related peptides: a multifunctional family of structurally related neuropeptides in insects. Adv Insect Physiol. 28, 267-329. (Doi 10.1016/S0065-2806(01)28012-6)

98 Elphick, M. R., Mirabeau, O. 2014 The Evolution and Variety of RFamide-Type Neuropeptides: Insights from Deuterostomian Invertebrates. Front Endocrinol (Lausanne). 5, 93. (10.3389/fendo.2014.00093)

99 Elphick, M. R., Semmens, D. C., Blowes, L. M., Levine, J., Lowe, C. J., Arnone, M. I., Clark, M. S. 2015 Reconstructing SALMFamide Neuropeptide Precursor Evolution in the Phylum Echinodermata: Ophiuroid and Crinoid Sequence Data Provide New Insights. Front Endocrinol (Lausanne). 6, 2. (10.3389/fendo.2015.00002)

100 Elphick, M. R. 2014 SALMFamide salmagundi: the biology of a neuropeptide family in echinoderms. Gen Comp Endocrinol. 205, 23-35. (10.1016/j.ygcen.2014.02.012)

101 Elphick, M. R., Price, D. A., Lee, T. D., Thorndyke, M. C. 1991 The SALMFamides: a new family of neuropeptides isolated from an echinoderm. Proc Biol Sci. 243, 121-127. (10.1098/rspb.1991.0020)

102 Fujisawa, Y., Kubota, I., Ikeda, T., Minakata, H., Muneoka, Y. 1991 A variety of Mytilus inhibitory peptides in the ABRM of Mytilus edulis: isolation and characterization. Comp Biochem Physiol C. 100, 525-531.

103 Bose, U., Suwansa-ard, S., Maikaeo, L., Motti, C. A., Hall, M. R., Cummins, S. F. 2017 Neuropeptides encoded within a neural transcriptome of the giant triton snail Charonia tritonis, a Crown-of-Thorns Starfish predator. Peptides. 98, 3-14. (10.1016/j.peptides.2017.01.004)

104 Zatylny-Gaudin, C., Cornet, V., Leduc, A., Zanuttini, B., Corre, E., Le Corguille, G., Bernay, B., Garderes, J., Kraut, A., Coute, Y., et al. 2016 Neuropeptidome of the Cephalopod Sepia officinalis: Identification, Tissue Mapping, and Expression Pattern of Neuropeptides and Neurohormones during Egg Laying. J Proteome Res. 15, 48-67. (10.1021/acs.jproteome.5b00463)

105 Woodhead, A. P., Khan, M. A., Stay, B., Tobe, S. S. 1994 Two new allatostatins from the brains of Diploptera punctata. Insect Biochem Mol Biol. 24, 257-263.

106 Woodhead, A. P., Stay, B., Seidel, S. L., Khan, M. A., Tobe, S. S. 1989 Primary structure of four allatostatins: neuropeptide inhibitors of juvenile hormone synthesis. Proc Natl Acad Sci U S A. 86, 5997-6001.

107 Pratt, G. E., Farnsworth, D. E., Fok, K. F., Siegel, N. R., McCormack, A. L., Shabanowitz, J., Hunt, D. F., Feyereisen, R. 1991 Identity of a second type of allatostatin from cockroach brains: an octadecapeptide amide with a tyrosine-rich address sequence. Proc Natl Acad Sci U S A. 88, 2412-2416.

108 Wegener, C., Gorbashov, A. 2008 Molecular evolution of neuropeptides in the genus Drosophila. Genome Biol. 9, R131. (10.1186/gb-2008-9-8-r131)

109 Peymen, K., Watteyne, J., Frooninckx, L., Schoofs, L., Beets, I. 2014 The FMRFamide-like peptide family in nematodes. Front Endocrinol. 5, (10.3389/fendo.2014.00090)

110 Martin-Duran, J. M., Pang, K., Børve, A., Le, H. S., Furu, A., Cannon, J. T., Jondelius, U., Hejnol, A. 2018 Convergent evolution of bilaterian nerve cords. Nature. 553, 45-50. (10.1038/nature25030)

111 Raikova, O. I., Reuter, M., Jondelius, U., Gustafsson, M. K. S. 2000 The brain of the Nemertodermatida (Platyhelminthes) as revealed by anti-5HT and anti-FMRFamide immunostainings. Tissue & Cell. 32, 358-365. (DOI 10.1054/tice.2000.0121)

